# Paracrine signalling between intestinal epithelial and tumour cells induces a regenerative programme

**DOI:** 10.1101/2020.12.16.423051

**Authors:** Guillaume Jacquemin, Annabelle Wurmser, Mathilde Huyghe, Wenjie Sun, Zeinab Homayed, Candice Merle, Meghan Perkins, Fairouz Qasrawi, Sophie Richon, Florent Dingli, Guillaume Arras, Damarys Loew, Danijela Vignjevic, Julie Pannequin, Silvia Fre

## Abstract

Tumours are complex ecosystems composed of different types of cells that communicate and influence each other. While the critical role of stromal cells in affecting tumour growth is well established, the impact of mutant cancer cells on healthy surrounding tissues remains poorly defined. Here, we uncovered a paracrine mechanism by which intestinal cancer cells reactivate foetal and regenerative Yap-associated transcriptional programs in neighbouring wildtype epithelial cells, rendering them adapted to thrive in the tumour context. We identified the glycoprotein Thrombospondin-1 (THBS1) as the essential factor that mediates non-cell autonomous morphological and transcriptional responses. Importantly, Thbs1 is associated with bad prognosis in several human cancers. This study reveals the THBS1-YAP axis as the mechanistic link mediating paracrine interactions between epithelial cells in intestinal tumours.

## Introduction

It is now well established that tumour formation and progression are vastly influenced by the crosstalk between cancer cells and their environment, involving complex remodelling of the extracellular matrix and interaction with stromal cells, such as cancer-associated fibroblasts, myofibroblasts, pericytes, vascular and lymphatic endothelial cells as well as different types of inflammatory immune cells (Marusyk *et al*, 2014; Cleary *et al*, 2014; Hanahan & Coussens, 2012). Remodelling of the tumour microenvironment has been shown to support tumour growth through neo-angiogenesis as well as via direct effects on cancer cells exposed to pro-inflammatory and pro-survival cytokines (Balkwill *et al*, 2012). However, paracrine interactions have mostly been studied among different cell types and little is known about communication between cancer and adjacent normal epithelial cells that could contribute to tumour formation and progression. Defining the mechanisms allowing paracrine interactions between tumour and normal epithelial cells requires an understanding of how different cells persist and expand within a tumour and is crucial to dissect intratumoral heterogeneity. It is noteworthy that these questions have lately received a special attention, and several studies addressing the co-existence and complex relationship between mutant and wildtype (WT) epithelial cells in the context of intestinal tumours have been published over the past year (Yum *et al*, 2021; Flanagan *et al*, 2021; van Neerven *et al*, 2021; Krotenberg Garcia *et al*, 2021).

We have recently reported that intestinal stem cells can be found within intestinal tumours and contribute to tumour growth (Mourao *et al*, 2019), suggesting the existence of bidirectional communications between tumour and normal epithelial cells. A recent study has also provided evidence that a parenchymal response of normal epithelial cells favours tumour growth and dissemination (Ombrato *et al*, 2019). The advent of 3D organotypic cultures able to faithfully recapitulate the morphology and physiology of intestinal cells in a mesenchyme-free environment has now allowed us to address the unresolved question of epithelial-specific interactions in the context of intestinal tumoroids.

Intestinal organoids are well-characterised stem-cell derived structures (Sato & Clevers, 2013). Importantly, organoids generated from normal mouse intestinal crypts consistently present a stereotypical “budding” morphology, with proliferative crypts (or buds) and a differentiated villus domain. On the other hand, cells derived from Apc mutant intestinal tumours generally grow as hyperproliferative and non-polarised hollow spheres or cysts (Drost *et al*, 2015; Sato *et al*, 2011; Jardé *et al*, 2013; Schwank *et al*, 2013; Germann *et al*, 2014; Onuma *et al*, 2013).

To study epithelial communications in a stroma-free environment, we analysed the influence of mutant organoids derived from primary mouse tumours (hereafter defined as “tumoroids”) on WT small intestinal organoids. We discovered that the co-culture of tumoroids and budding organoids quickly induced a hyperproliferative cystic morphology (referred to as “cysts” hereafter) in a fraction of WT organoids. This interaction did not require cell contact, as the effect was recapitulated by the conditioned medium from tumoroids. We found that the secreted glycoprotein Thrombospondin-1 (THBS1) was responsible for mediating these paracrine communications, through Yap pathway activation. Under the influence of tumour-derived THBS1, WT cells activate the YAP signalling pathway and induce foetal and regenerative transcriptional programs, which cause their hyperproliferation and failure to properly differentiate. Importantly, we show that the THBS1/YAP1 signalling axis we discovered in organoids is conserved in both mouse and human colon cancer and propose that this early mechanism of non-cell autonomous epithelial communication is critical for the establishment of a primary tumour. Of medical relevance, we also found that THBS1 expression is necessary for tumoroids growth, providing a potential therapeutic target. These studies offer novel insights into the molecular mechanisms responsible for tumour growth and provide an attractive therapeutic avenue in targeting THBS1 to reduce tumour complexity and heterogeneity.

## Results

### Tumour cells induce a cancer-like behaviour in wildtype epithelial intestinal cells

To mimic intra-tumoral heterogeneity in stroma-free conditions, we co-cultured tumoroids derived from primary intestinal tumours of Apc^1638N/+^ mutant mice (Fodde *et al*, 1994), labelled by membrane tdTomato (Muzumdar *et al*, 2007) with WT GFP-marked small intestinal organoids, derived from lifeact-GFP mice (Riedl *et al*, 2008) (Fig. EV1A). Within 24-48 hours of co-culture with tumoroids, WT organoids (up to 20%), normally displaying the stereotypical budding morphology (Sato *et al*, 2009) (Fig. 1A), adopted an unpolarised hollow cystic shape presenting a diameter larger than 100µm (indicated by arrows in Figure 1B), closely resembling the morphology of Apc mutant tumoroids (Schwank *et al*, 2013) (Fig. 1C). To assess if this morphological change was due to mid-range paracrine or juxtacrine signals, we cultured WT organoids in conditioned medium (cM) from either wildtype (WT-cM) or tumour (T-cM) organoids. Consistent with our observations from co-cultures, WT organoids exposed to T-cM (Fig. 1E), but not to WT-cM (Fig. 1D), grew as cysts, suggesting that tumoroids secrete factors able to morphologically alter WT epithelial cells. Tumoroids derived from different primary tumours reproducibly induced the cystic “transformation”, albeit to variable extents (Fig. EV1B). Of interest, the conditioned medium from Apc^-/-^ organoids (derived from VillinCre^ERT2^; Apc^flox/flox^ mice), where Apc knock-out was induced by Cre recombination and did not rely on spontaneous Apc LOH, reproduced the effect of T-cM and induced a cystic morphology in WT organoids, confirming a direct effect caused by aberrant Wnt signalling (Fig. EV1C). The morphological change was all the more remarkable as it occurred within 6 to 12 hours of exposure to T-cM (Fig. EV1D-E) and was reversed after 2-4 days, if no fresh medium was added, suggesting exhaustion of the responsible factor(s) and ruling out the possibility of acquired genetic mutations in normal organoids. Since a cystic organoid morphology has been linked to Wnt pathway activation, we analysed the expression of the quantitative Wnt reporter 7TG (Brugmann *et al*, 2007). As shown in Figure 1F, cystic organoids grown in T-cM did not present canonical Wnt pathway activation (Fig. 1F right panel), unlike organoids stimulated with the small molecule CHIR99021, an inhibitor of the enzyme GSK-3, widely used to simulate Wnt activation (Ring *et al*, 2003) (Fig. 1F middle panel). Confirming these observations, we could not observe any significant difference in the number of Lgr5+ cells in the presence of T-cM, compared to both WT-cM and normal ENR (Fig. EV1F), whereas exposure to the GSK3 inhibitor CHIR99021 (ENRC) resulted, as expected, in a significant increase of GFP+ cells. Despite absence of Wnt activation and similar numbers of Lgr5-expressing cells (Fig. 1F and Fig. EV1F), we found that T-cM exposed cystic organoids presented a higher proportion of cycling cells (compare cystic “C” and budding “B” in Fig. 1G and EV1G). Moreover, while proliferative cells were restricted to the crypts in control organoids exposed to WT-cM (Fig. 1H), cystic organoids in T-cM displayed undifferentiated proliferative cells scattered throughout the newly formed cysts (Fig. 1I). The increase in proliferative cells is linked to defective enterocyte differentiation, as shown by loss of Keratin 20 expression (Fig. 1I).

**Fig. 1.**
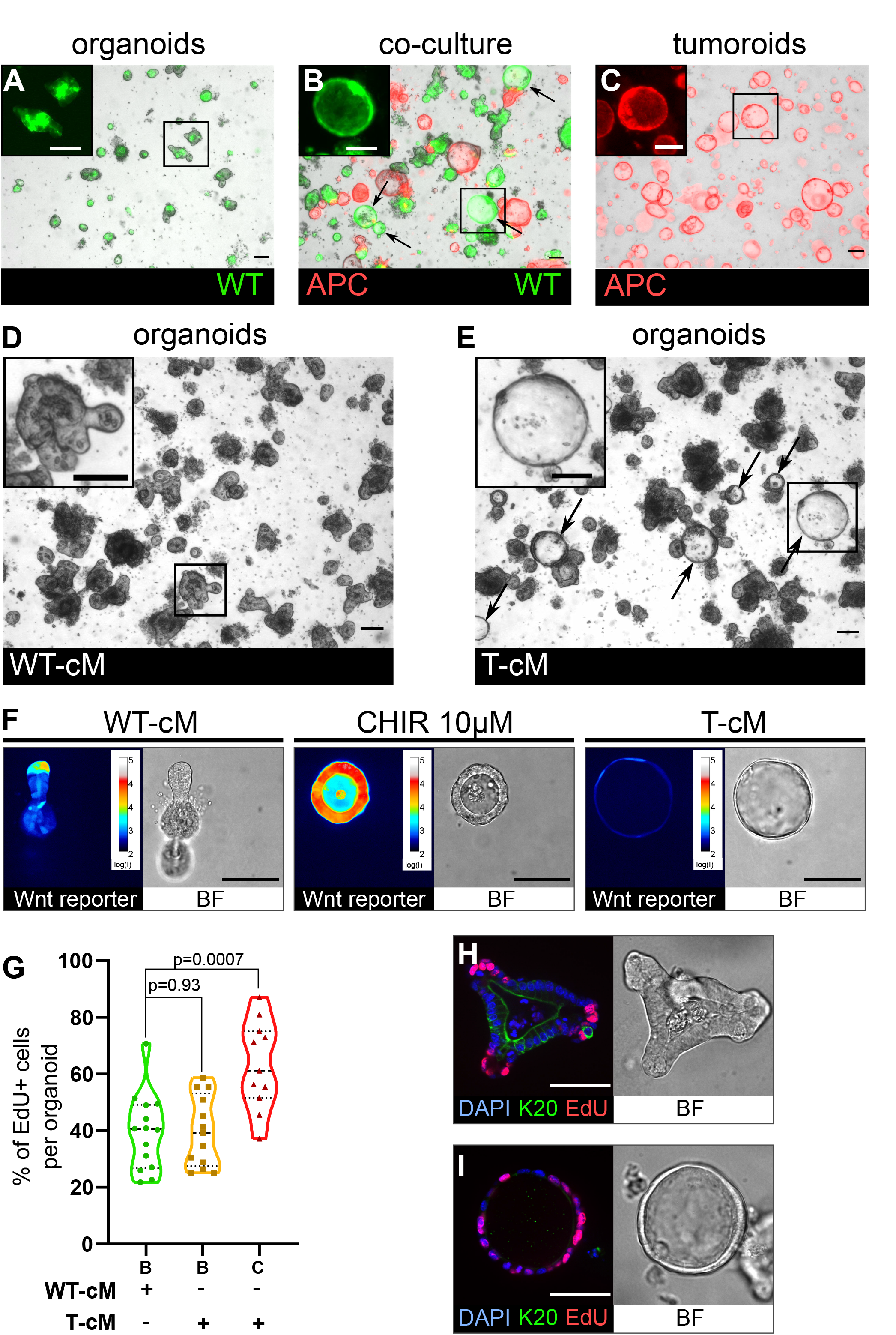
Tumoroids secrete soluble factors that induce a tumour-like cystic morphology in WT organoids. (A) WT budding organoids marked by lifeact-GFP (in green) after 24h in culture in organoid medium (ENR). (B) WT organoids marked by lifeact-GFP after 24h in co-culture with tdTomato-expressing tumoroids in ENR. Arrows indicate WT (green) cystic organoids. (C) APC mutant cystic tumoroids marked by tdTomato after 24h in culture. (D-E) WT budding organoids cultured for 24h in conditioned medium from WT organoids (WT-cM in D) or from tumoroids (T-cM in E). Arrows indicate WT cystic organoids in E. (F) Representative images of WT organoids expressing the Wnt reporter 7TG exposed to WT-cM, 10µM CHIR99021 (CHIR 10 µM) or T-cM for 24h. Pseudo-colour shows log10 intensities of the reporter fluorescence. (G) Quantification of the percentage of EdU+ cells per organoid for budding organoids grown in WT-cM (B – WT-cM, n=14), budding organoids grown in T-cM (B – T-cM, n=13) or cystic organoids grown in T-cM (C – T-cM, n=9). (H-I) Immunofluorescence for proliferative cells (EdU in red) and differentiated cells (anti-Keratin 20 in green) in WT organoids grown in WT-cM (H) or in T- cM (I) for 24h. The corresponding bright field images (BF) are shown in the right panels. DAPI stains DNA in blue. Scale bar = 100µm. Statistical analysis was performed with two-tailed unpaired Welch’s t-tests.

### Thrombospondin-1 (Thbs1) mediates the organoid morphological and behavioural change

Cancer cells are known to secrete numerous factors to remodel the tumour microenvironment. Having established that the morphological change was mediated by proteins present in the T-cM, since the effect was abolished upon proteinase K treatment (Fig. EV2A), we performed a quantitative proteomics analysis by SILAC (Stable Isotope Labelled Amino acids in Culture) mass spectrometry to define the composition of the T-cM relative to WT-cM. Gene Ontology analysis of the proteins over-represented in T-cM showed enrichment in cell adhesion, wound healing and generally ECM-related GO terms (Fig. EV2C). We selected secreted factors that were enriched in T-cM compared to WT-cM (Fig. EV2B) and explored their possible involvement in the morphological “transformation” using neutralising antibodies. Among the tested candidates, we found that neutralisation of the secreted glycoprotein Thrombospondin-1 (THBS1) alone was sufficient to entirely abolish the morphological change of WT organoids after 24h (Fig. 2A and EV2D). Neutralisation of Thbs1 with three blocking antibodies, targeting different epitopes of the protein to exclude any potential non-specific binding, was sufficient to completely block the cystic morphology (Fig. 2B, EV2E). These experiments demonstrated that THBS1 was necessary for the observed cystic phenotype.

**Fig. 2.**
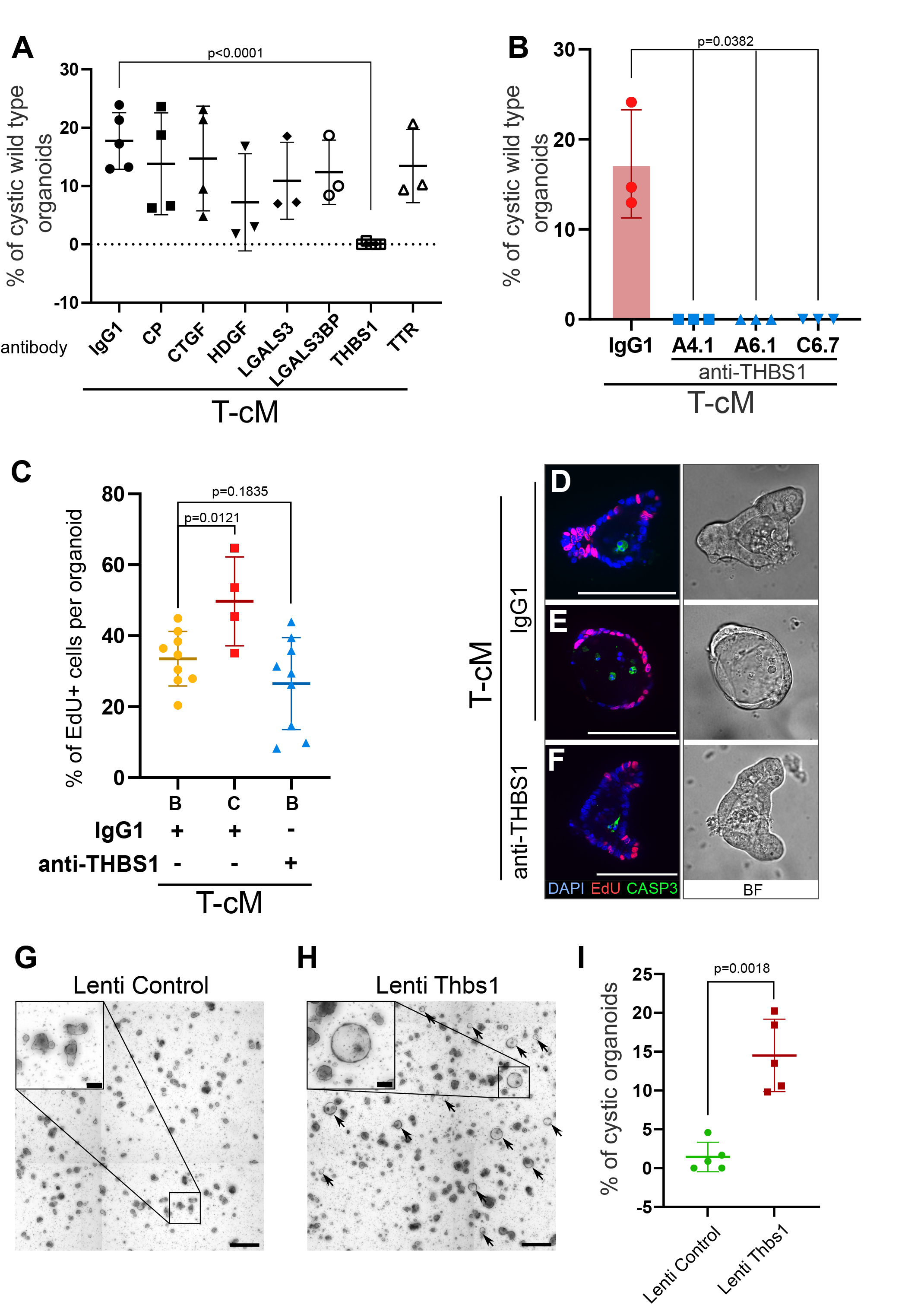
Thrombospondin-1 is necessary and sufficient for the morphological “transformation” of WT organoids. (A) Quantification of the percentage of WT cystic organoids in T-cM upon neutralisation with blocking antibodies against Ceruloplasmin (CP), Connective tissue growth factor (CTGF), Hepatoma derived growth factor (HDGF), Galectin-3 (LGALS3), Galectin-3 binding protein (LGALS3BP), Thrombospondin-1 (THBS1), Transthyretin (TTR) (5 µg/ml). (B) Quantification of the percentage of WT cystic organoids in T-cM upon neutralisation with three different blocking antibodies against THBS1 (clones A4.1, A6.1 and C6.7 at 5 µg/ml). (C) Quantification of the percentage of EdU+ cells (2h pulse) per organoid for WT budding (B – IgG1, n=9) or cystic organoids (C – IgG1, n=4) exposed to T-cM in the presence of IgG1 or antibodies anti-THBS1 (B – anti-THBS1, n=9). (D-F) Whole-mount immunostaining for proliferation (EdU in red) and apoptosis (anti-cleaved caspase-3, CASP3 in green) of WT organoids exposed to T-cM with anti-IgG1 control (D-E) or anti-THBS1 (F) antibodies. DAPI stains DNA in blue. The corresponding bright field (BF) images are shown on the right panels. (G-H) Representative pictures of self-transformed WT organoids overexpressing Thbs1 (Lenti-Thbs1 in H) and control organoids infected with an empty vector (Lenti-Control in G). Black arrows indicate cystic organoids. (I) Quantification of the percentage of cystic organoids in Thbs1- expressing cultures (Lenti-Thbs1) versus control cultures (Lenti-Control) grown for 24h in ENR medium (n=5). Scale bars = 100µm in D-F and in the insets of G-H and 500µm in G-H low magnification. Graphs indicate average values ± SD. Statistical analysis was performed with two- tailed unpaired Welch’s t-tests.

In order to test if THBS1 neutralisation was also able to rescue the ectopic proliferation, we counted the proliferative cells per organoid. Consistent with our previous observations, cystic organoids presented ectopically proliferating cells when cultured with IgG1 control antibodies (Fig. 2C, E). However, addition of anti-THBS1 antibodies abolished ectopic proliferation and restricted EdU+ cells exclusively to the crypts (Fig. 2C, F). Treatment with anti-THBS1 did not present toxicity to normal organoids, as no increase in apoptotic cells was observed (Fig. 2D-F). Moreover, Thbs1 was sufficient to induce the morphological change, since its ectopic expression in WT organoids (Fig. EV2F) also led to cyst development (Fig. 2G-I) to the same extent as T-cM (Fig. 2I). Of note, culture of WT organoids in the presence of recombinant Thbs1 did not elicit any effect, possibly due to the lack of essential post-translational modifications.

### Thrombospondin-1 is necessary for the growth of tumoroids but not of normal organoids

We observed that Thbs1 is exclusively expressed by tumours but not by normal intestinal cells (Fig. EV2B, EV5A-C); surprisingly, we found that THBS1 is also essential for tumoroids growth. Indeed, neutralisation of THBS1 for 48h specifically reduced tumoroid survival and considerably arrested their growth (Fig. 3A-D, G-J, EV3A). Importantly, the same tumoroid growth inhibition was observed upon Thbs1 genetic deletion (Fig. 3E-F) using CRISPR-Cas9 knock-out (Fig. EV3B-E). Quantification of the proportion of dividing cells showed that neutralisation of THBS1 significantly reduced tumoroids’ proliferative capacity (Fig. 3M-N, P), without affecting the growth of normal organoids (Fig. 3K-L, O) suggesting a promising therapeutic avenue.

**Fig. 3.**
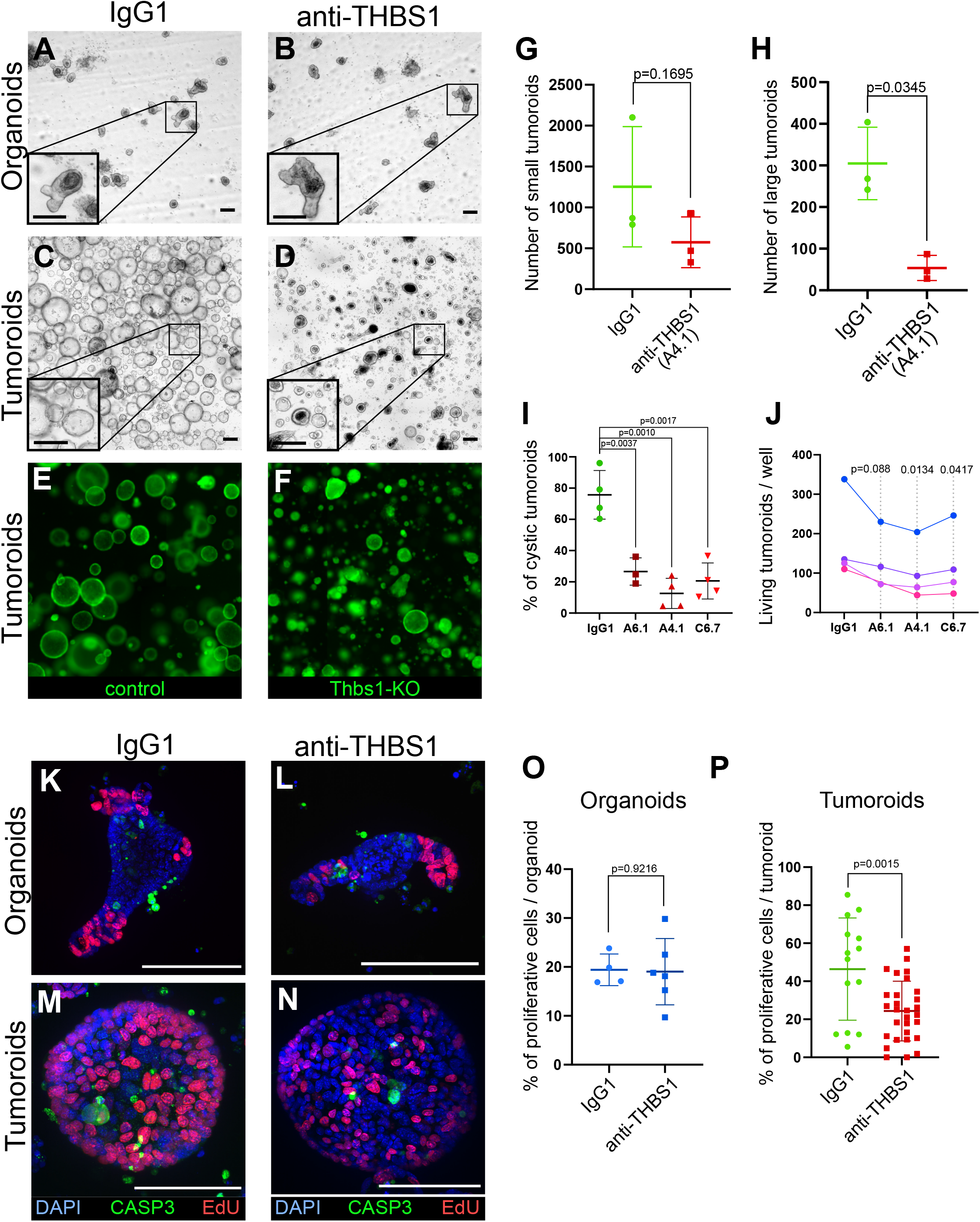
Thrombospondin-1 is essential for the growth of tumoroids. (A-D) Representative bright field images of WT organoids (A-B) or tumoroids (C-D) incubated with IgG1 isotype control antibodies (A and C) or anti-THBS1 A6.1 neutralising antibody (B and D) (10 µg/ml). (E-F) Representative images of tumoroids infected with a lentivirus CRISPR-GFP without sgRNA (control in E) or with a sgRNA targeting Thbs1 (Thbs1-KO in F) 48h after replacement of single cell seeding medium (ENRC) by tumoroid medium (EN). (G-H) Quantification of the number of tumoroids upon antibody neutralisation relative to their size: small tumoroids between 30 to 150µm in G; large tumoroids of more than 150 µm diameter in H. (I) Quantification of the percentage of cystic tumoroids upon treatment by IgG1 isotype control antibodies or 3 different neutralising antibodies targeting THBS1 (as indicated) for 48h. (J) Paired quantification of the number of living tumoroids derived from 4 independent tumours (from 4 mice) upon treatment with three neutralising antibodies targeting THBS1 for 48h. Antibody concentration: 10µg/ml. (K-N) Immunofluorescence staining for proliferative cells (EdU in red) and apoptosis (anti-cleaved Caspase-3, CASP3 in green) in WT organoids (K-L) or tumoroids (M-N) exposed to IgG1 isotype control antibodies (K and M) or to anti-THBS1 A6.1 neutralising antibody (L and N). (O-P) Quantification of EdU+ cells per organoid (O) or tumoroid (P) in the presence of IgG1 control or anti-THBS1 (A6.1) antibodies. Scale bars = 100µm. Graphs indicate average values ± SD. Statistical analysis was performed with paired Student’s t-test in G, H, I and J and with two-tailed unpaired Welch’s t-tests in O and P.

### WT organoids activate a regenerative/foetal transcriptional program upon the influence of tumoroids conditioned medium

In order to decipher the molecular responses of WT epithelial cells to T-cM, we obtained the gene expression profiles of WT organoids exposed to either T-cM or WT-cM, along with the transcriptional signature of the tumoroids from which the corresponding T-cM was derived. Interestingly, we found that T-cM induced transcriptional responses enriched for genes upregulated in cancer, including colorectal adenoma (in red in Fig. EV4A), suggesting that, alongside the typical tumour-like cystic morphology, WT cells also acquire signatures characteristics of tumour cells when exposed to T-cM. Furthermore, this analysis revealed that organoids grown in the presence of T-cM showed enrichment of genes of the Yes-associated protein (YAP)/Hippo pathway (Fig. EV4A in green). Given the intricate relationship between Wnt signalling and YAP in the intestine, suggesting that tumour formation requires additional signals other than Wnt, that induce YAP nuclear translocation (Cai *et al*, 2015; Azzolin *et al*, 2014; Gregorieff *et al*, 2015; Taniguchi *et al*, 2015, 2017; Guillermin *et al*, 2021), we assessed the involvement of the YAP pathway in T-cM mediated phenotypes. First, we compared our RNA sequencing results to the YAP activation signature from intestinal organoids (Gregorieff *et al*, 2015) using Gene Set Enrichment Analysis (GSEA), and found a strong correlation in both WT organoids (Fig. 4A) and tumoroids (Fig. 4D). Consistent with recent studies describing YAP activation as an integral part of the regenerative and foetal programs of the normal intestinal epithelium (Yui *et al*, 2018), we found a robust association with the reported physiological “foetal human colitis” intestinal signature (Yui *et al*, 2018) (Fig. 4B,E), suggesting a link between the morphological change we characterised and reactivation of regenerative/foetal programs occurring during tumorigenesis. These findings indicate that WT organoids in the presence of tumour secreted factors, including THBS1, switch their transcriptional program from a Wnt-dependent homeostatic to a Wnt-independent, YAP-dependent regenerative/foetal-like response, repressing differentiation genes (Fig. 1I and 4C, F) without significantly affecting Wnt signalling (Fig. 1F and EV4B).

**Fig. 4.**
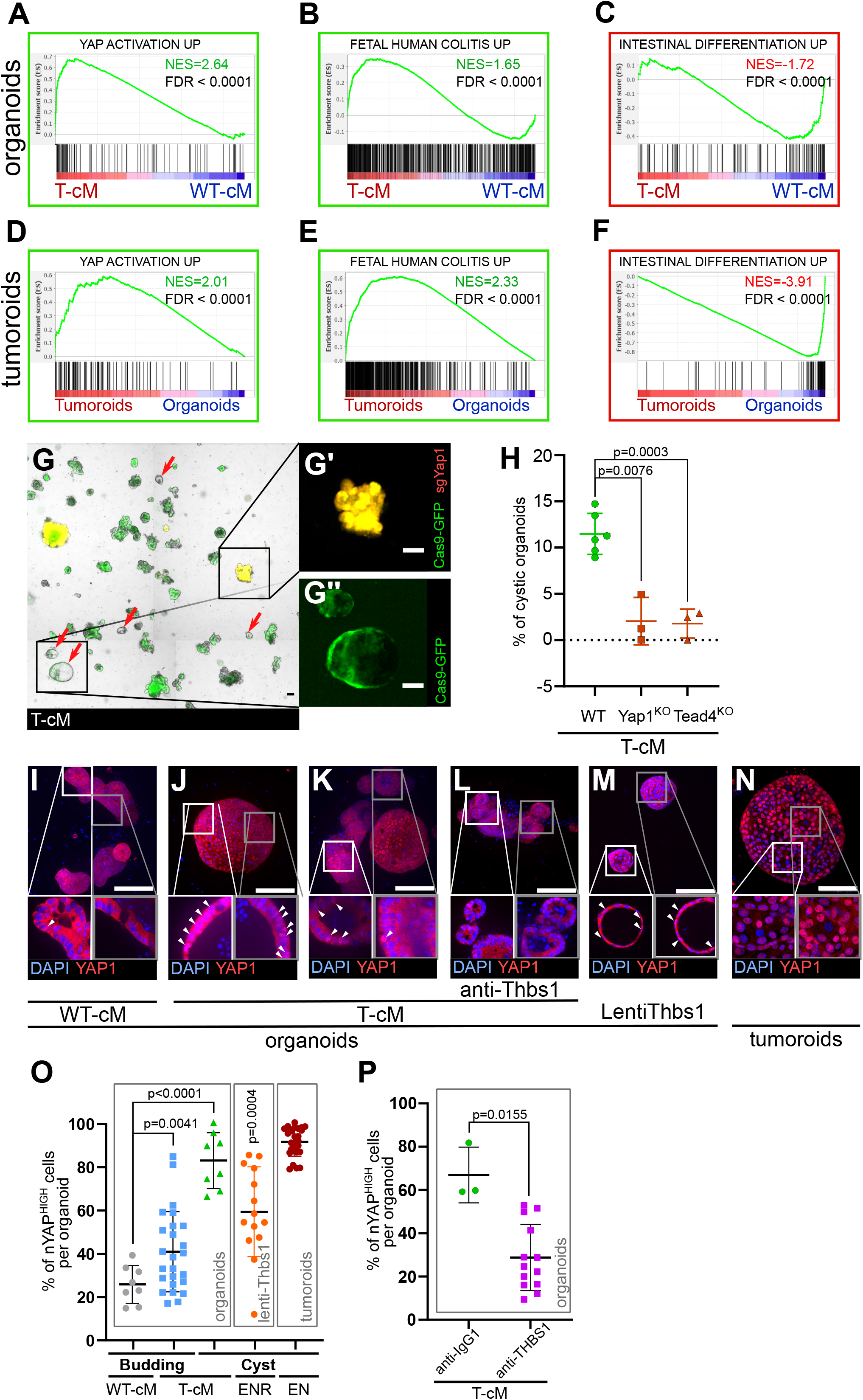
T-cM induces YAP pathway activation and a foetal-like state in WT organoids. Gene Set Enrichment Analyses (GSEA) showing the correlation between differentially expressed genes in WT organoids cultured in T-cM (A-C) or in tumoroids (D-F) and the indicated transcriptional signatures. NES: Normalised Enrichment Score; green NES = positive correlation; red NES = inverse correlation. (G) Representative image of WT organoids expressing Cas9-GFP (in green) transduced with a sgRNA targeting Yap1 (sgYap1 in red). Higher magnification of a budding Yap1^KO^ organoid (in yellow in G’) and a cystic Yap1^WT^ organoid expressing only Cas9-GFP but no sgRNA (in green in G’’). (H) Percentage of cystic organoids induced by exposure to T-cM in WT, Yap1^KO^ or Tead4^KO^ organoids, as indicated. (I-N) Max projections of immunostaining for YAP1 (in red) of WT organoids exposed to WT-cM (I), or T-cM (J-L) for 24h presenting cystic (J) or budding (K and L) morphologies. Organoids in (L) are treated by neutralising antibodies targeting THBS1 (A6.1), which rescues the budding morphology. Organoids in (M) overexpress THBS1 (LentiThbs1) and tumoroids are shown in (N). DAPI stains DNA in blue. White arrowheads pinpoint YAP^HIGH^ cells in Z-section insets. (O) Quantification of the percentage of nuclear YAP (nYAP^HIGH^) cells/organoid based on the ratio of nuclear vs. cytoplasmic YAP1 in cystic and budding WT organoids grown in WT-cM or T-cM for 24h, in WT organoids overexpressing Thbs1 (lenti-Thbs1) or in tumoroids, as indicated. (P) Quantification of the percentage of YAP^HIGH^ cells/organoid based on the ratio of nuclear vs. cytoplasmic YAP1 immunofluorescence in WT organoids cultured with T-cM and control IgG1 or anti-THBS1 (A6.1) antibodies for 24h. Scale bars correspond to 100µm in G, I-N. Graphs indicate average values ± SD. Statistical analysis was performed with two-tailed unpaired Welch’s t-tests. For the lenti- Thbs1 sample, a Welch’s corrected t-test was applied to compare the percentage of nYAP^HIGH^ cells/organoid between Thbs1-expressing organoids and WT organoids infected with an empty lentivirus.

To assess the functional significance of YAP pathway activation, we pharmacologically blocked it using Verteporfin, an inhibitor of the YAP-TEAD interaction (Liu-Chittenden *et al*, 2012), and observed a complete loss of cystic organoids with no discernible effects on their growth (Fig. EV4C-D). We further corroborated these results through the genetic deletion by CRISPR/Cas9 of Yap1 and one of its cellular effectors, the transcription factor Tead4 (Guillermin *et al*, 2021), found upregulated upon exposure to T-cM (Fig. EV4E-F). Yap1 or Tead4 knock-out in WT organoids (Fig. 4G) caused a considerable decrease in the proportion of cystic organoids induced by T-cM, suggesting that the YAP/Hippo pathway mediates the morphological change (Fig. 4H). We further found that WT organoids, upon T-cM exposure, displayed a higher number of cells with nuclear YAP1, a readout of YAP pathway activation and a typical characteristic of tumoroids (Fig. 4I-K, O). Notably, T-cM induces nuclear YAP accumulation in both cystic and budding organoids (Fig. 4J, K, O), suggesting that YAP activation is necessary but not sufficient to induce the switch to the cystic phenotype.

Importantly, neutralisation of THBS1 with three blocking antibodies was sufficient to rescue both the cystic phenotype and the percentage of cells presenting YAP nuclear accumulation, which dropped to control levels (Fig. 4L, P). Corroborating the key role of THBS1 in YAP activation, we also found that lentiviral overexpression of THBS1 (Lenti-Thbs1) was sufficient to trigger both cystic shapes and YAP nuclear translocation (Fig. 4M, O). Furthermore, we assayed the effect of T-cM onto organoids derived from mouse colon (colonoids) and confirmed that T-cM also induced YAP activation and promoted proliferation in colonoids (Fig. EV4G-J), as was the case for small intestinal organoids, consolidating the relevance of our findings to colon cancer.

### Thbs1-expressing and YAP-activated tumour cells are mutually exclusive in mouse tumours

To further substantiate the *in vivo* relevance of our results, we induced acute Apc loss in VillinCre^ERT2^;Apc^flox/flox^ mice for a short time (4 days) and found that Thbs1 was ectopically expressed by Apc mutant intestinal epithelial cells, indicating that Thbs1 expression is induced by Wnt activation (Fig EV5A, B). To demonstrate that YAP nuclear accumulation is induced in WT epithelial cells neighbouring mutant tumour cells, we induced mosaic Apc loss in both VillinCreERT2;Apc^flox/+^ and VillinCreERT2;Apc^flox/flox^ mice, allowing us to study non-recombined WT epithelial cells adjacent to or within Apc mutant tumours. These experiments showed that in both Apc heterozygotes (Fig. 5F) and homozygotes (Fig. 5G) mice, the majority of the cells presenting nuclear YAP do not coincide with Apc mutant cells (displaying high levels of cytoplasmic and nuclear beta-catenin, indicative of Wnt activation), but they are always in close proximity to the mutant cells.

**Fig. 5.**
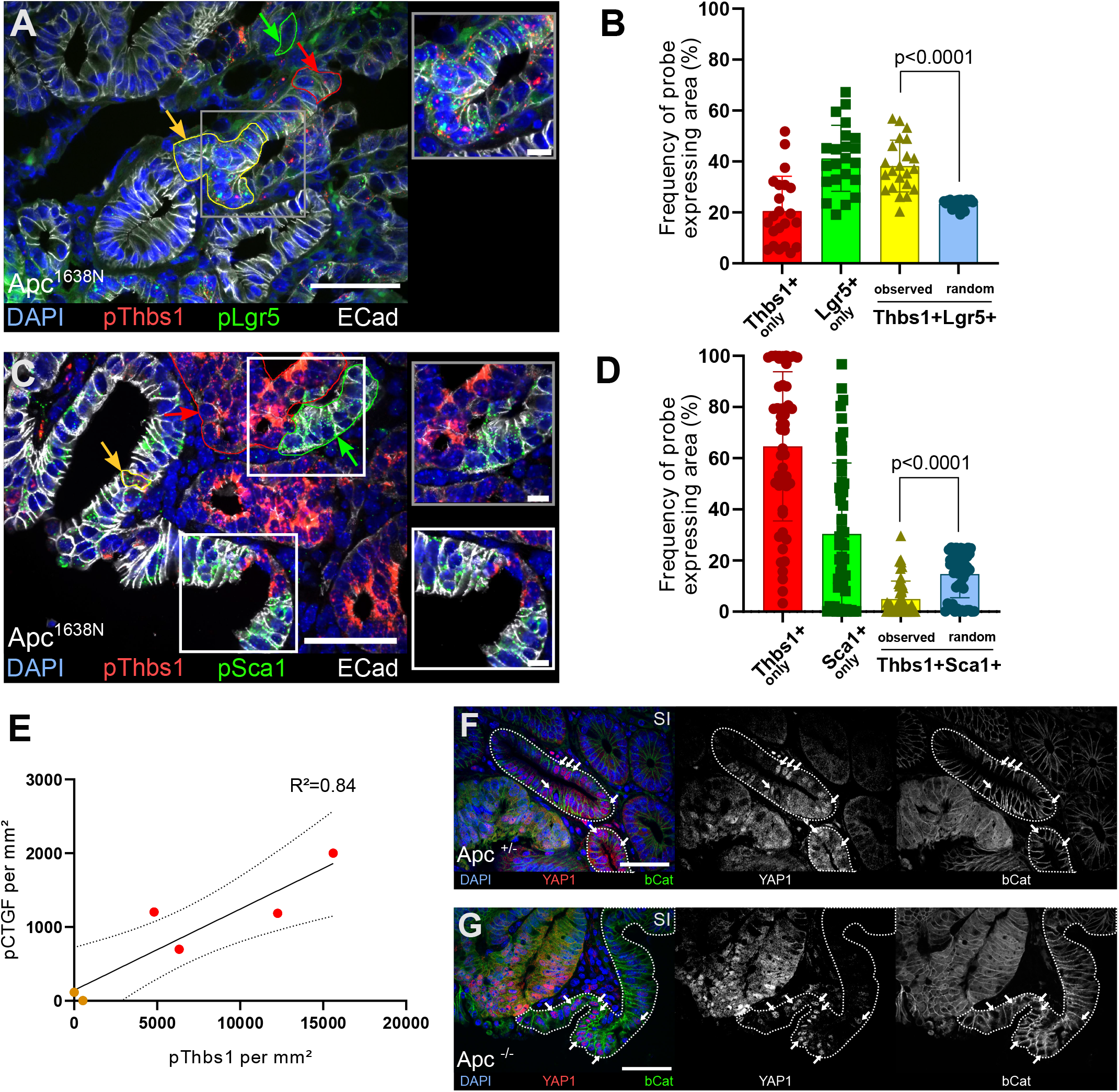
Thbs1 is expressed by Lgr5+ cancer stem cells *in vivo* and induces YAP activation in neighbouring epithelial cells. (A, C) Representative section of Apc mutant intestinal tumours analysed by smFISH for Thbs1 (pThbs1, red dots) and Lgr5 (pLgr5, green dots in A) or the YAP target Sca1 (pSca1, green dots in C). Examples of segmented and processed ROI that were automatically counted as co-localisation (Thbs1+/Lgr5+ in A or Thbs1+/Sca1+ cells in C outlined in yellow and indicated by yellow arrows) or single probe expression (outlined in red or green and indicated by arrows of the corresponding colour) are shown. E-cadherin demarcates epithelial cells in white and DAPI labels nuclei in blue in A and C. (B, D) Quantification of the frequency of tumour regions expressing exclusively one probe or co-expressing two probes (yellow): Thbs1 only in red or Lgr5 only in green (B); Thbs1 only in red or Sca1 only in green (D). The observed frequencies of co-localisation (yellow in B) or mutual exclusion (yellow in D) are statistically significant compared to the calculated probability of random co-expression (blue columns) (n=22 sections from 2 tumours in B and n=51 sections from 5 tumours in D). (E) Correlation of the number of RNA molecules (dots/mm²) detected by smRNA FISH for the YAP target CTGF and Thbs1 in mouse intestinal tumours. Red dots indicate large tumours (≥8mm), orange dots small tumours (<8mm). Dashed lines indicate 95% confidence intervals. (F-G) Representative sections of tumours derived from villinCre;Apc^flox/+^ (Apc^+/-^ in F) or villinCre;Apc^flox/flox^ (Apc^-/-^ in G) immunostained for YAP1 (in red) and beta-catenin (in green). WT glands displaying membrane-bound beta-catenin, adjacent to mutant areas presenting diffuse cytoplasmic/nuclear beta-catenin expression are demarcated by dashed lines. White arrows indicate examples of cells showing high levels of nuclear YAP. Scale bars = 50µm and 10µm in insets. Statistical analysis was performed with Wilcoxon test in B-D and linear regression test with 95% confidency in E.

Consistent with these findings, in mouse tumours, we found that Thbs1+ cells largely coincide with the cells expressing the widely accepted Wnt target genes Lgr5 (Barker *et al*, 2009) (Fig. 5A, B, EV5E) and Axin2 (Fig. EV5D) while they do not express the differentiation marker Keratin 20 (Fig. EV5F). The RNA probe recognizing Thbs1 co-localizes with the THBS1 protein visualized by antibody staining (Fig. EV4G). Furthermore, single molecule RNA FISH showed a clear and highly significant mutual exclusion between Thbs1-expressing cells and cells showing YAP activation, as assessed by expression of the YAP targets Sca1 (Fig. 5C, D, Fig. EV5J), Ctgf (Fig. EV4H) and Cyr61 (Fig. EV4I), indicating the presence of two distinct tumour cell populations: one Thbs1+/Sca1- (64.61% ±29.14%) and one Thbs1-/Sca1+ (30.40% ±27.72%) (Fig. 5D). Of relevance to colon cancer, these results are confirmed both in small intestinal adenomas (Apc^1638N^ in Fig. 5 and Fig. EV5A-K) and in chemically-induced colon tumours (Fig EV5L, M). At the whole-tumour scale, Thbs1 and CTGF expression are highly correlated (Fig. 5E, R²=0.84).

### The THBS1-YAP axis is conserved in early stages of human colorectal cancer

To establish if the mechanism mediating paracrine cellular communication that we uncovered is conserved in human colon cancer, we then analysed a cohort of 10 human colon tumours (5 low grade adenomas and 5 invasive carcinomas) for their expression of THBS1, LGR5 and YAP (Fig. 6C-D, 6E-F). The main components of the signalling axis we have uncovered, Thbs1, Ctgf and Cyr61, but not the Wnt target gene Lgr5, are highly correlated in bulk transcriptomics of human colorectal samples (Fig. 6A). Supporting our results in mouse adenomas, the analysis of human tumours at different stages revealed that Thbs1 is highly expressed in Lgr5+ cells only in early-stage adenomas (Fig. 6C) but not in carcinomas (Fig. 6D). High co-expression of Thbs1 and Lgr5 in adenomas is accompanied by the presence of extensive tumour regions rich in cells presenting nuclear YAP (Fig. 6E), which were not visible in invasive adenocarcinomas (Fig. 6F). This intriguing observation can explain why no significant correlation between Thbs1 and Lgr5 expression was found in our in-silico analysis of human advanced colon cancer (Fig. 6A). These results, combined with our findings in organoids and transgenic mice, suggest a key role of the Thbs1-YAP axis in tumour initiation. Of further prognostic relevance, Thbs1 expression (Fig. 6G), as well as the levels of the YAP target gene Ctgf (Fig. 6H), are strongly correlated with poor prognosis in human colon cancer and also in kidney and gastric tumours.

**Fig 6.**
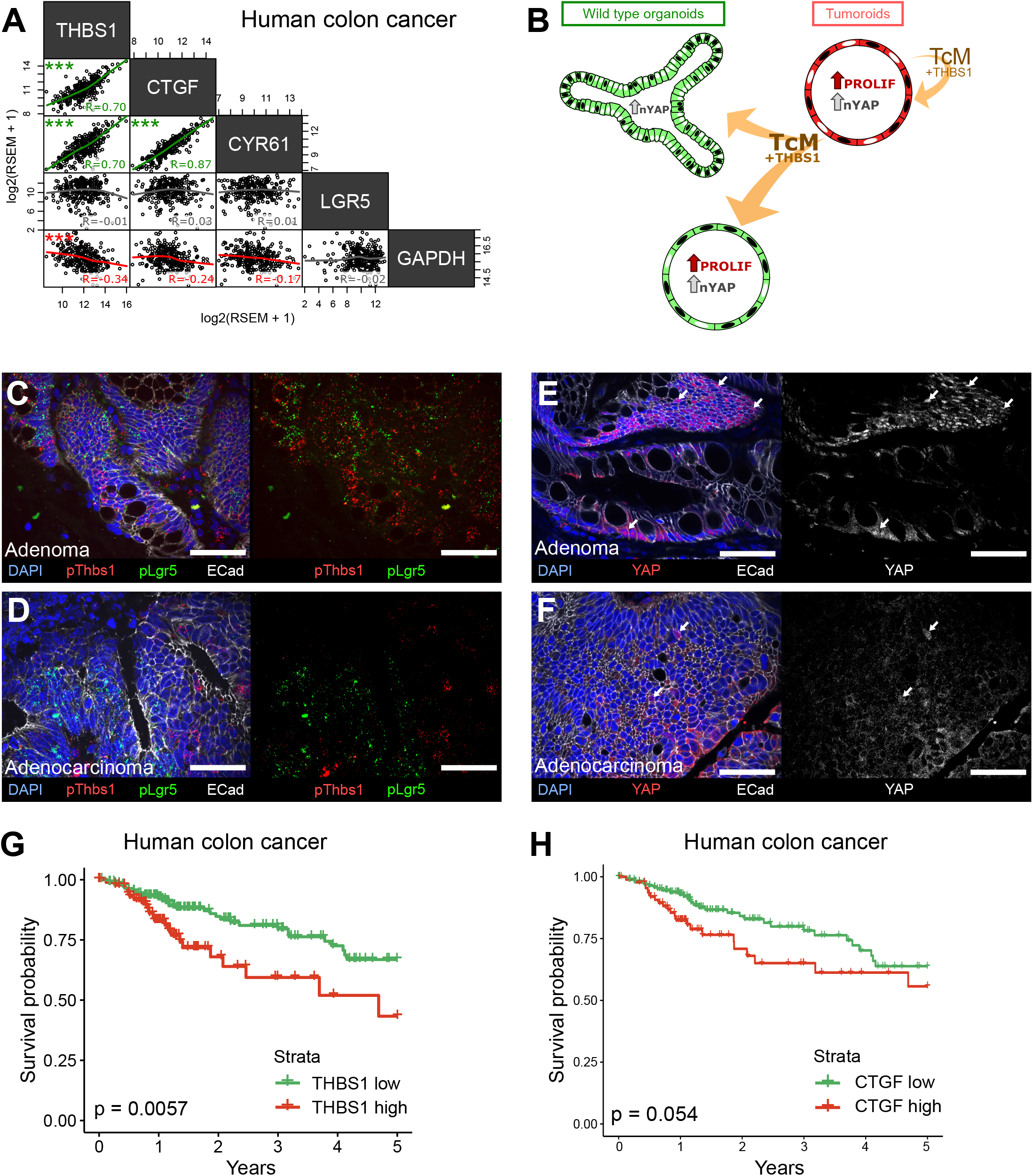
The THBS1-YAP pathway operates in human low grade adenomas. (A) Correlation matrix between the expression levels of THBS1and the YAP targets CTGF, CYR61 and LGR5 in human colon tumours from the TCGA colon cancer bulk datasets. R indicates Spearman coefficient. (B) Graphical summary of paracrine interactions between WT organoids and tumoroids along the THBS1-YAP axis. Mutant tumoroids “corrupt” genetically WT organoids by secreting THBS-1 (orange arrows). This results in YAP1 nuclear translocation (black nuclei in organoids or tumoroids) and ectopic proliferation as well as cystic morphology in a subset of organoids. (C-F) Representative sections of low grade human adenomas (C, E) or advanced human carcinomas (D, F) processed by smRNA FISH for Thbs1 (pThbs1, red dots) and Lgr5 (pLgr5, green dots in C-D) or immunostained with anti-YAP1 antibodies (E-F). White arrows highlight tumour cells presenting high nuclear YAP. n = 5 human low grade adenomas in C, E and n = 5 advanced human adenocarcinomas in D, F. (G-H) Overall survival of colon cancer patients stratified into THBS1 high (red) and low (green) groups (in G) or CTGF high (red) and low (green) groups (in H) using a threshold of the top one-third quantile of gene expression.

## Discussion

Our results implicate that both in intestinal tumours derived from spontaneous Apc loss and in chemically-induced colonic tumours, cancer cells can directly recruit surrounding epithelial cells through Wnt-driven expression and secretion of the glycoprotein THBS1, which results in aberrant activation of a regenerative/foetal transcriptional program mediated by the YAP pathway (Fig. 6B), a driver of intestinal regeneration and tumorigenesis (Gregorieff *et al*, 2015). THBS1 is overexpressed in a large number of solid tumours, but its role in cancer is controversial. Constitutive deletion of Thbs1 in Apc^Min/+^ mice led to an increase in the number and aggressiveness of tumours, which was interpreted as a consequence of its anti-angiogenic role (Gutierrez *et al*, 2003). However, THBS1 has been reported to promote the attachment of cells to the extracellular matrix, favouring cancer cell migration and invasion (Sid *et al*, 2008; Tuszynski *et al*, 1987). Indeed, a study using a model for inflammation-induced colon carcinogenesis (AOM/DSS) in Thbs1^-/-^ mice showed a 5-fold reduction in tumour burden, suggesting a role for THBS1 in tumour progression (Lopez-Dee *et al*, 2015). These contradictory results are most likely due to the multifaceted effects of THBS1, depending on which cells secrete it and which cells respond. Of interest, a recent study proposed that THBS1 induces focal adhesions (FAs) and nuclear YAP translocation through interaction with αvβ1 integrins in the aorta (Yamashiro *et al*, 2019). Moreover, FAs have been shown to directly drive YAP1 nuclear translocation in the foetal intestine and upon inflammation in adult colon (Yui *et al*, 2018).

Here, we found that cancer cell-derived THBS1 can “corrupt” WT epithelial cells in organoids, independently of stroma-derived effects, allowing us to address the specific role of THBS1 on epithelial cells. Surprisingly, we found that three neutralising antibodies targeting different epitopes of THBS1 were all able to block its effect on WT organoids. These results may indicate that THBS1 neutralisation is not due to block of a specific ligand-receptor interaction but rather to the steric interference with the trimerization of the large soluble THBS1 isoform (450kDa) that may no longer be free to diffuse through the Matrigel. Alternatively, it is possible that that several THBS1 domains are involved in YAP activation, consistent with reports indicating that different THBS1 domains interact with integrins (Resovi *et al*, 2014).

Our results, corroborated by consistent observations in mouse and human intestinal tumours, uncovered a novel function for the secreted multidomain glycoprotein THBS1 in affecting the behaviour of normal epithelial cells surrounding a nascent tumour. The coexistence of WT and mutant cells within nascent tumours has been the subject of recent interest and, consistent with our results, paracrine communication between epithelial cells has been found to Induce YAP pathway activation (Flanagan *et al*, 2021; Yum *et al*, 2021; van Neerven *et al*, 2021; Krotenberg Garcia *et al*, 2021). However, it is still debated whether YAP has a tumour suppressor (Barry *et al*, 2013; Cheung *et al*, 2020) or oncogenic role (Zanconato *et al*, 2016). Indeed, while the recent studies suggest that cancer cells actively eliminate WT cells by cell competition, facilitating tumour expansion, our results indicate that the recruitment of WT cells by cancer cells happens at the very early steps of tumour formation and may be required for the cancer cells to seed within the hyperplastic epithelium. However, consistent with the recent literature and our analysis of human tumours, at later stages of tumorigenesis, the fitter mutant cells outcompete WT cells, as shown by the decrease in cells expressing both THBS1 and nuclear YAP in advanced human adenocarcinomas (Fig. 6C-F). We thus believe that the observed differences in cell behaviour could depend on the kinetics of the effects of tumour-secreted factors on WT cells, distinguishing very early responses (within 24-48 hours) analysed in this study and later outcomes (van Neerven *et al*, 2021). Also, only some WT cells, responding to tumour-secreted factors by activating YAP, may be able to survive within tumours, while the majority of WT cells would be outcompeted, as proposed by (Krotenberg Garcia *et al*, 2021).

Based on the results we report here, we propose a model for tumour initiation where mutant cells can “corrupt” surrounding WT epithelial cells by secreting THBS1, leading to YAP activation in the receiving cells. Through the paracrine mechanism we unravelled, causing reactivation of a regenerative foetal-like transcriptional program, WT epithelial cells initially thrive in nascent tumours. In such a scenario, normal cells within tumours, which would escape specific therapies targeting mutant cells, could be identified by their hallmark of YAP activation, providing a novel diagnostic tool. Of relevance, THBS1 neutralisation showed a tumour-specific toxicity; we thus propose that THBS1 may represent a therapeutic target for colon cancer, potentially applicable to other epithelial tumours.

## Materials and Methods

### Ethics Statement

All studies and procedures involving animals were in strict accordance with the recommendations of the European Community (2010/63/UE) for the Protection of Vertebrate Animals used for Experimental and other Scientific Purposes. The project was specifically approved by the ethics committee of the Institut Curie CEEA-IC #118 and approved by the French Ministry of Research with the reference #04240.03. We comply with internationally established principles of replacement, reduction, and refinement in accordance with the Guide for the Care and Use of Laboratory Animals (NRC 2011). Husbandry, supply of animals, as well as maintenance and care of the animals in the Animal Facility of Institut Curie (facility license #C75–05–18) before and during experiments fully satisfied the animal’s needs and welfare. Suffering of the animals has been kept to a minimum; no procedures inflicting pain have been performed.

### Statistics and Reproducibility

Experiments were performed in biological and technical replicates as stated. For each experiment, we have used at least n=3 organoid lines originating from n=3 different mice, and experiments with at least n=3 replicates were used to calculate the statistical value of each analysis. All graphs show mean ± SD. Statistical analysis was performed with two-tailed unpaired Welch’s t-tests, unless otherwise stated.

### Transgenic mouse models

All mouse lines used have been previously described. Intestinal tumours were generated in Apc1638N mutant mice (Fodde *et al*, 1994), crossed to the R26^mTmG^ line (Muzumdar *et al*, 2007) or in villinCreERT2 (el Marjou *et al*, 2004) crossed to Apc Delta14 (Colnot *et al*, 2004) mice and Lgr5-GFP (Barker *et al*, 2007) mice were kindly provided by H. Clevers. GFP expressing wildtype organoids were generated from the LifeAct-GFP mouse line (Riedl *et al*, 2008). Organoids used for KO experiments were obtained by crossing R26-LSL-Cas9-GFP (Platt *et al*, 2014) and R26CreERT2 mouse lines (Ventura *et al*, 2007). All mice used were of mixed genetic background.

### Chemically induced colon tumour model

AOM/DSS colon carcinogenesis experimental protocol. To induce colon tumours, we followed the protocol from Tanaka and colleagues (Tanaka *et al*, 2003): Notch1-Cre^ERT2^/R26^mTmG^ mice of 5-7 months of age received a single intraperitoneal injection of Azoxymethane (AOM, Sigma #A5486) followed by Dextran Sulfate Sodium (DSS, MP Biomedicals #160110) administration (3% in drinking water) one day after the AOM injection for 5 consecutive days. General health status and mouse body weight were monitored daily during and after treatment. To verify the presence of colon tumours, 2 mice were checked 1 month after the first cycle of DSS treatment, but no tumours were detected (only signs of inflammation). We administered another cycle of DSS (3% in drinking water) for 3 days and tumour formation was monitored by colonoscopy using a Karl-Storz endoscopic system.

### Human tumours

Five low grade adenomas and five invasive adenocarcinomas were obtained from the Centre of Biological Resources of Institut Curie and examined by the service of pathology.

### Organoids cultures

*Wild type organoids:* Wildtype organoids were cultured and passaged as previously described (Sato *et al*, 2009) and derived from the small intestine or colon of 2-5 months old mice. The Matrigel-crypts mix was plated as 50µl drops in 24-well plates or 35 µl drops in 8-well Ibidi imaging chambers (Ibidi 80827) for whole mount staining. After polymerisation the Matrigel drop was covered with wild type organoid medium containing EGF, Noggin, R-spondin1 (ENR) for small intestinal organoids or EGF, Noggin, R-spondin1, CHIR99021, Y27632, Wnt3a (ENRCYW) for colonoids. Reagents, media and buffers are listed in Supplementary Table S1. Factors, inhibitors and neutralising antibodies that were added to the medium are indicated in Supplementary Table S2.

*Tumoroids:* Apc1638N heterozygous mice of more than 6 months of age were dissected and intestinal tumours were harvested using forceps and micro-dissection scissors to reduce contamination with adjacent healthy tissue. Periampullary tumours were excluded from the study to avoid contamination with stomach cells. To remove the remaining healthy tissue surrounding the extracted tumour, tumours were incubated in 2 mM EDTA in PBS (pH=8.0) for 30 minutes at 4°C. Tumours were then briefly vortexed to detach the remaining normal tissue, leaving clean spheres. In order to dissociate tumour cells, the tumour was chopped into 1-3 mm fragments using a razor blade and digested in 66% TrypLE (Thermofischer 12605010) diluted in PBS, for 10 min at 37°C under continuous agitation at 180 RPM. The supernatant containing the dissociated cells was harvested and fresh 66% TrypLE was added to the remaining fragments for another 10 minutes. The supernatant was strained using a 70 µm cell strainer. Cells were then centrifuged at 400g for 5 minutes at 4°C, suspended in DMEM-F12 (2% Penicillin Streptomycin) and plated in 50% Matrigel drops as described for wildtype organoid cultures. After polymerisation, 300 µl of EN medium containing EGF and Noggin (EN) was added. The medium was replaced every 1-2 weeks. Tumoroids were passaged every 1 (for line expansion) or 3 to 4 weeks (for medium conditioning). Reagents, media and buffers are listed in Supplementary Table S1. The medium composition is summarised in Supplementary Table S2.

### Co-culture assay

Wildtype and tumour organoid cultures were started at least two weeks before co-culture in order to use stable and exponentially growing cultures. These passages guaranteed morphologically homogeneous organoids. After passage of wildtype and tumour organoids, the fragments were mixed at approximately 3:1 wildtype organoid to tumoroids ratio. Mixed fragments were plated as described above. After Matrigel polymerisation at 37°C, ENR medium (300 µl/well) was added. Reagents, media and buffers are listed in Supplementary Table S1.

### Conditioned medium assay

WT organoids were passaged as previously described. After Matrigel polymerisation, 150 µl of ENR 2X concentrated and 150 µl of conditioned medium (cM) were added. Reagents, media and buffers are listed in Supplementary Table S1. Analyses were performed between 24 and 48h after plating, unless otherwise specified.

#### Tumoroid conditioned medium preparation

Established tumoroid cultures (more than two passages) were grown for 1 week. After one week of expansion, fresh medium was added and conditioned for 1 to 2 weeks depending of the organoid density. Immediately after harvesting, the conditioned medium was centrifuged at 400g for 5 minutes at 4°C to remove big debris and cell contamination. The supernatant was recovered and centrifuged again at 2.000g for 20 minutes at 4°C and then ultra-centrifuged at 200.000g for 1.5 hours at 4°C in order to remove extracellular vesicles. The supernatant was frozen in liquid nitrogen and stored at -80°C.

#### WT conditioned medium preparation

Established WT organoid cultures were passaged and grown for 3 days in order to obtain high density cultures. Then, fresh ENR medium was conditioned for a maximum of 1 week and prepared as described above.

### Organoid freezing

Matrigel drops containing organoids in exponential growth were collected in PBS and centrifuged twice at 400g for 5 minutes at 4°C to remove Matrigel and debris. The organoids were suspended at a high density in Cryostor10 freezing medium (six wells of organoids per 1 ml of cryogenic medium). Organoids were incubated for 10 min in Cryostor10 before being frozen. Reagents, media and buffers are listed in Supplementary Table S1.

### Organoid thawing

Organoids were quickly thawed at 37°C and suspended in 5ml of FBS prior to centrifugation at 400g for 5 minutes at 4°C. The organoids were suspended in DMEM-F12 with 2% Penicillin/Streptomycin and mixed with Matrigel at a 1:1 ratio as described above. After polymerisation at 37°C, ENR or EN medium was added. Due to the FBS impact on organoid morphology, the thawed organoids were passaged at least once and carefully checked for their morphology prior to use. Reagents, media and buffers are listed in Supplementary Table S1.

### Organoid infection

Wild type or tumour organoid cultures were started at least a week before transduction. To plate 8 wells of infected organoids, 12 wells of exponentially growing organoids were harvested in cold Cell Recovery Solution and incubated on ice for 15 minutes to dissolve the Matrigel. Organoids were then centrifuged at 400g for 5 minutes at 4°C. For cell dissociation, the pellet was suspended in 2 ml of AccuMax (SIGMA A7089) containing CHIR99021, Y27632 (CY) and incubated at 37°C for 8 minutes. Digestion was then stopped by adding 2 ml of DMEM-F12 containing B27 and CY. Organoids where further mechanically dissociated by pipetting up and down 40-50 times. The cell suspension was then centrifuged at 400g for 5 minutes at 4°C. Organoids were carefully suspended in 300 µl of 66x concentrated virus (see below) containing CY and Transdux reagents prior to addition of 300 µl of cold-liquid Matrigel and plated as previously described. After polymerisation at 37°C, 300 µl/well of ENR-CY medium was added. ENR-CY was replaced by ENR two days later. This is essential to avoid a morphological change due to exposure to CHIR99021. After 4 days, small organoids were passaged and cultured following the protocol described above. Reagents, media and buffers are listed in Supplementary Table S1.

### Lentiviruses

#### Plasmids

Lenti-7TG was a gift from Roel Nusse (Addgene plasmid # 24314; http://n2t.net/addgene:24314; RRID: Addgene_24314). The Lenti-sgRNA-mTomato (LRT) construct was obtained by replacement of the GFP sequence with tandem Tomato in the Lenti-sgRNA-GFP (LRG), a gift from Christopher Vakoc (Addgene plasmid # 65656; http://n2t.net/addgene:65656; RRID: Addgene_65656). LentiCRISPRv2GFP was a gift from David Feldser (Addgene plasmid # 82416; http://n2t.net/addgene:82416; RRID: Addgene_82416). LentiThbs1Tg is a lentiORF expressing mouse Thbs1 (NM_011580) –myc-DKK (Origene #MR211744L3V). pMD2.G was a gift from Didier Trono (Addgene plasmid # 12259; http://n2t.net/addgene:12259; RRID: Addgene_12259). psPAX2 was a gift from Didier Trono (Addgene plasmid # 12260; http://n2t.net/addgene:12260; RRID: Addgene_12260).

#### sgRNA cloning

Both lentivirus backbones used for the knock-out experiments harboured the GeCKO cloning adaptors (Shalem *et al*, 2014; Sanjana *et al*, 2014). sgRNA inserted in either vectors are listed in Supplementary Table S3.

#### Virus production

Lentiviral particles were produced in HEK 293T cells. Day 0: 8 x 10^6^ cells were plated onto a T75 flask in 10 ml of complete medium (see Supplementary Table S1). Day 1: cells were transfected using PEI/NaCl. For this, two solutions were prepared: mix A contained 625µl NaCl 150mM + 75µl PEI and mix B contained 625µl NaCl 150mM + 6µg of plasmid DNA at a molar ratio of 4:3:2 (lentiviral vector: psPAX2: pMD2.G). Mix A and B were incubated for 5 min at room temperature and then mixed and incubated for 15 min at RT before being added drop by drop on top of the cells. Medium was changed at day 2 and 10ml of supernatant containing virus particles was collected at day 3 and 4. The supernatants were centrifuged for 5 min at 1500rpm at 4°C in order to remove dead cells and debris. The virus particles were then concentrated to 300µl using Amicon Ultra Centrifugal Filters (UFC910024 SIGMA) by spinning at 1000g at 4°C for 1h. *sgRNA validation* sgRNAs cut efficiency was assessed in Mouse Embryonic Fibroblasts (MEFs) infected with a lentivirus expressing the Cas9 enzyme along with blasticidin-resistance (Addgene plasmid #52962). 48 hours upon infection, infected MEFs were selected in 10µg/ml Basticidin for 7 days. Short guide RNAs (sgRNAs) targeting Thbs1, Yap1 and Tead4 (Table S3) were cloned in the LRT lentiviral vector (expressing red Tomato fluorescent protein) and produced as described above. After transduction of the MEFs-Cas9 with LRT-Thbs1, Yap1 or Tead4, cells were FACS-sorted based on their red fluorescence and their genomic DNA extracted. For each sgRNA, the cut site region (±200bp) was PCR amplified and sequenced using the same primers (Table S3). Chromatograms were manually analysed using ApE (v2.0.61) to confirm the precise cut site, which induced mutations starting at -3bp before the PAM sequence.

### Mass Spectrometry

#### Sample preparation

Isotope labelled conditioned medium requires pre-loading of isotopic amino acids in order to detect all proteins synthetized and secreted by cells (Ong *et al*, 2002). To label newly produced proteins, two essentials isotopically labelled amino acids (IAA), arginine and lysine, were added to the culture medium. WT organoids were labelled with [²H_4_]-lysine (Lys4) and [^13^C_6_]-arginine (Arg6) for 2 weeks (2 passages), whilst tumoroids were labelled with [^13^C_6_ N_2_]-lysine (Lys8) and of [^13^C_6_ N_4_]-arginine (Arg10) for 2 weeks (2 passages). Then, medium was conditioned as described above using SILAC-ENR (2x1 week) or SILAC-EN (1x2weeks). After a functional assay confirming their transforming capacities, conditioned media were concentrated. Subsequently, 2 ml of tumoroids conditioned media or 4 ml of wild type conditioned media were precipitated by cold acetone. Dried protein pellets were then recovered with 50 µl of 2X Laemli buffer with SDS and β−mercapto-ethanol (0.1%), boiled at 95°C for 5 minutes and centrifuged 5 min at 14000g.

#### MS Sample processing

Gel-based samples were cut in 8 bands and in-gel digested as described in standard protocols. Briefly, following the SDS-PAGE and washing of the excised gel slices proteins were reduced by adding 10 mM Dithiothreitol (Sigma Aldrich) prior to alkylation with 55 mM iodoacetamide (Sigma Aldrich). After washing and shrinking of the gel pieces with 100% acetonitrile, trypsin / LysC (Promega) was added and proteins were digested overnight in 25 mM ammonium bicarbonate at 30°C. Extracted peptides were dried in a vacuum concentrator at room temperature and re-dissolved in solvent A (2% MeCN, 0.3% TFA) before LC-MS/MS analysis.

#### LC-MS/MS analysis

Liquid chromatography (LC) was performed with an RSLCnano system (Ultimate 3000, Thermo Scientific) coupled online to an Orbitrap Fusion Tribrid mass spectrometer (MS, Thermo Scientific). Peptides were trapped on a C18 column (75 μm inner diameter × 2 cm; nanoViper Acclaim PepMapTM 100, Thermo Scientific) with buffer A’ (2/98 MeCN/H2O in 0.1% formic acid) at a flow rate of 2.5 µL/min over 4 min. Separation was performed on a 50 cm x 75 μm C18 column (nanoViper Acclaim PepMapTM RSLC, 2 μm, 100Å, Thermo Scientific) regulated to a temperature of 55°C with 1) a linear gradient of 5% to 30% buffer B (100% MeCN in 0.1% formic acid) at a flow rate of 300 nL/min over 100 min. Full-scan MS was acquired in the Orbitrap analyser with a resolution set to 120,000, a mass range of m/z 400–1500 and a 4 × 105 ion count target. Tandem MS was performed by isolation at 1.6 Th with the quadrupole, HCD fragmentation with normalised collision energy of 28, and rapid scan MS analysis in the ion trap. The MS2 ion count target was set to 2 × 104 and only those precursors with charge state from 2 to 7 were sampled for MS2 acquisition. The instrument was run at maximum speed mode with 3 s cycles.

#### Data Processing

Data were acquired using the Xcalibur software (v 3.0) and the resulting spectra were interrogated by SequestHT through Thermo Scientific Proteome Discoverer (v 2.1) with the Mus musculus Swissprot database (022017 containing 16837 sequences and 244 common contaminants). The mass tolerances in MS and MS/MS were set to 10 ppm and 0.6 Da, respectively. We set carbamidomethyl cysteine, oxidation of methionine, N-terminal acetylation, heavy ^13^C_6_ N_2_- Lysine (Lys8) and ^13^C_6_ N_4_-Arginine (Arg10) and medium ^2^H_4_-Lysine (Lys4) and C_6_-Arginine (Arg6) as variable modifications. We set specificity of Trypsin digestion and allowed two missed cleavage sites. The resulting files were further processed by using myProMS (v 3.5)(Poullet *et al*, 2007). The SequestHT target and decoy search results were validated at 1% false discovery rate (FDR) with Percolator. For SILAC-based protein quantification, peptides XICs (Extracted Ion Chromatograms) were retrieved from Thermo Scientific Proteome Discoverer. Global MAD normalization was applied on the total signal to correct the XICs for each biological replicate (n=3). Protein ratios were computed as the geometrical mean of related peptides. To estimate ratio significance, a t-test was performed with the R package limma (Ritchie *et al*, 2015) and the false discovery rate was controlled using the Benjamini-Hochberg procedure (Benjamini & Hochberg, 1995) with a threshold set to 0.05. Proteins with at least two peptides, a 2-fold enrichment and an adjusted p-value < 0.05 were considered significantly enriched in sample comparisons.

#### Pathway enrichment

Gene ontology (GO) terms enrichment analysis used the proteins significantly enriched in sample comparisons (T-cM/WT-cM; 2peptides, fold change >2, adjusted p-value<0.05) and the unique proteins to T-cM. GO biological processes, cellular components, and molecular functions were analysed using the UniProt-GOA Mouse file (v. 20181203). Significant GO terms had a P < 0.05.

### Immunofluorescence (IF)

#### Organoids staining

For whole mount IF, organoids were grown on 8well-chamber slides (Ibidi 80827). For EdU staining, a 2h pulse of EdU (10µM, Carbosynth Limited NE08701) preceded fixation. After fixation using 4% Paraformaldehyde (Euromedex 15710) in PBS for 1h at room temperature, organoids were washed with PBS and permeabilised in PBS + 1% triton X-100 (Euromedex 2000-C) for 1h at room temperature. Organoids were then incubated with 150µl of diluted antibodies (see Supplementary Table S4) in blocking buffer (PBS, 2% BSA, 5% FBS, 0.3% Triton X-100) overnight at room temperature. After 3 washes of 5 min in PBS, 150µl of secondary antibodies were added together with DAPI diluted in PBS and incubated at room temperature for 5h. Organoids were then washed with PBS for 3 times for 5 min each and stored in 1:1 ratio PBS and glycerol (Euromedex 15710) before imaging. For EDU staining, EDU signal was revealed after the secondary antibody step, using the EDU click-it kit according to the manufacturer’s protocol (Thermofisher scientific C10340).

#### Human sections IF staining

Paraffin sections 3µm of human samples were deparaffinised and rehydrated using the standard protocol of xylene/ethanol gradient. Antigens were unmasked by boiling the slides in a citrate-based solution (Eurobio-Abcys H-3300). Slides were then incubated in blocking buffer (PBS, 2% BSA, 5% FBS). Antibodies were then added in blocking buffer (see Supplementary Table S4) overnight at 4°C. After 3 washes of 5 min in PBS, secondary antibodies were added together with DAPI diluted in PBS and incubated at room temperature for 2h. Slides were mounted in Aqua- poly/mount (Tebu Bio 18606-5).

#### Mouse sections IF staining

Intestinal tissue samples were fixed overnight at room temperature with 10% formalin prior to the paraffin embedding. 4um FFPE sections were prepared for beta-catenin / YAP co-staining. Briefly, the tissue sections were deparaffinized 5 times in xylene 5 mins each, rehydrated 5 times in ethanol 100% 5 mins each then once in ethanol 70% for 10 mins. A heat mediating antigen retrieval was made using boiling Sodium citrate tribasic dihydrate solution 10mM, PH=6 (Sigma S4641) for 20 mins. The sections were then blocked and permeabilized with 5% Donkey serum 0.01% Triton X-100 for 30 min at RT before being incubated with rabbit anti-YAP dilution 1/100 (Signalling Technology #14074) and mouse anti beta-catenin (BD 610153) (dilution 1/200) overnight at 4°C. The next day, slides were washed 3 times in PBS Tween20 0.01% and then incubated for 1h at room temperature with matching secondary antibodies donkey anti-rabbit A594 (Jackson Research 711-546-152) and donkey anti-mouse A488 (Jackson Research 715-546-150) (dilution 1/500) with DAPI. Slides were mounted using Fluoromount Aqueous Mounting Medium (Sigma F4680).

### Single molecule RNA Fluorescence In Situ Hybridization (smRNA FISH)

smRNA FISH was performed on mouse tissue cryosections or human paraffin-embedded tumour sections using RNAscope® Multiplex Fluorescent Detection Kit v2 kit (ACD 323110) and pipeline following manufacturer’s recommendations. Thbs1 mRNA were labelled using RNAscope® Probe- Mm-Thbs1-C3 (#457891-C3) or Hs-THBS1-C2 (#426581-C2), CTGF mRNA were labelled using RNAscope® Probe- Mm-CTGF (#314541) and Lgr5 mRNA were labelled using RNAscope® Probe- Mm-Lgr5 (#312171) or Hs-LGR5-C3 (#311021-C3). In order to subsequently perform immunostaining after the FISH, a protease III step not exceeding 20 minutes was included. Subsequent antibody staining was performed as described.

### Epithelial masks generation

To quantify RNAscope results only in epithelial cells, epithelial masks were generated using E-cadherin immunostaining. After Ilastik training allowing segmentation of E-cadherin-stained membranes, masks were smoothed by closing function (iteration=15) and holes filling. Generated masks were manually corrected for consistency.

### smRNA FISH dots quantification

Raw images were segmented using Ilastik (v1.3.2) and training on negative controls (background), positive controls and experimental slides (dots). Segmented masks were cleaned using FIJI through an opening function, 2px Gaussian blur and Moments threshold. Generated masks were manually checked for consistency with raw data. Aggregates of dots were partially rescued using watershed function and dots (size=2-250 circularity=0.50-1.00) analysed using build-in Analyse Particle function. Epithelial dots were obtained by multiplication of dot masks by the corresponding epithelial masks previously generated.

### Image acquisition

Images were obtained on an Inverted Wide Confocal Spinning Disk microscope (Leica) using 40x/1.3 OIL DIC H/N2 PL FLUOR or 20x/0.75 Multi immersion DIC N2 objectives and Hamamtsu Orca Flash 4.0 camera. Images were captured using Metamorph. Whole plate acquisition was performed using a dissecting microscope and Cell Discoverer 7 (Leica). Images were captured with ZEN. For RNAscope experiments, images were obtained on PLAN APO 40x/1.3NA o objective on an upright spinning disk (CSU-X1 scan-head from Yokogawa) microscope (Carl Zeiss, Roper Scientific France), equipped with a CoolSnap HQ2 CCD camera (Photometrics). Images were captured using Metamorph.

### Nuclear/cytoplasmic quantification

Nuclear IF ratios were obtained using a custom-made ImageJ macro. This macro segmented the nuclei on the DAPI channel using Otsu Threshold, Watershed and Particles analysis. For each segmented nucleus, a cytoplasmic halo of 8 pixels was generated and excluded from the DAPI mask to avoid false cytoplasmic measure in neighbouring nuclei. The mean intensities of the segmented nucleus and cytoplasmic regions of interest (nROI and cROI) were then measured in the IF channel (nIF and cIF). Cells were counted only if: area ratio nucleus/cytoplasm > 0.5, avoiding bias of pixel sampling either due to miss-segmentation or to overcrowded regions. Results (nIF, cIF and ratio) were then processed on Microsoft Excel. A density curve of the nIF / cIF ratio was performed for each category of organoid in order to observe peaks trends of positive and negative nuclei for IF. Threshold was defined manually in the inter-peak region at 1.1 (YAP) and 1.25 (EdU). To exclude low or non-specific signal, a minimal mean intensity cut-off for cIF was established from the experimental images. Using the threshold and the cut-off, the percentage of nIF^HIGH^ cells per organoid was calculated.

### RNA-sequencing

#### Sample preparation

Organoids were harvested using Cell recovery solution as described above. After 15 minutes of Matrigel dissolution, organoids were pelleted at 500g for 5 minutes. Pellets were recovered in 1ml of PBS in 1.5ml tubes and pelleted again at same conditions. RNA extraction was performed using RNEasy Mini Kit (Qiagen) following manufacturer’s recommendations. Total RNA integrity (RINe) were subjected to quality control and quantification using an Agilent TapeStation instrument showing excellent integrity (RNA Integration Number, RIN=10). Nanodrop spectrophotometer was used to assess purity based on absorbance ratios (260/280 and 260/230).

#### RNA Sequencing

RNA sequencing libraries were prepared from 1µg of total RNA using the Illumina TruSeq Stranded mRNA Library preparation kit which allows to prepare libraries for strand specific mRNA sequencing. A first step of polyA selection using magnetic beads was performed to address sequencing specifically on polyadenylated transcripts. After fragmentation, cDNA synthesis was performed followed by dA-tailing before ligation of the TruSeq indexed adapters (Unique Dual Indexing strategy). PCR amplification was finally achieved to create the final cDNA library. After qPCR quantification, sequencing was carried out using 2x100 cycles (paired-end reads, 100 nucleotides) on an Illumina NovaSeq 6000 system (S1 flow cells) to get around 45M paired-end reads per sample. Fastq files were generated from raw sequencing data using bcl2fastq where demultiplexing was performed according to indexes.

#### RNA-seq data processing

Sequencing reads were aligned on the Mouse reference genome (mm10) using the STAR mapper (v2.5.3a) (Dobin *et al*, 2013). Protein-coding genes from the Gencode annotation (vM13) have been used to generate the raw count table. Overall sequencing quality controls report a very high sequencing quality, a high fraction of mapped reads, and a high enrichment in exonic reads.

#### Differential analysis

Expressed genes (TPM>=1 in at least one sample) have then been selected for supervised analysis. The raw count table was normalised using the TMM method from the edgeR R package (v3.25.9)(Robinson *et al*, 2010), and the limma (Ritchie *et al*, 2015) voom (v3.39.19) functions were applied to detect genes with differential expression. In order to compare tumoroids versus wiltype samples, we design a linear model as follow:

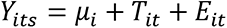

where T is the type effect (T={WT-cM, T-cM, Tumoroids}). We then restricted the dataset to WT-cM and T-cM samples, and applied the following model:

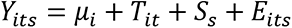

where T is the type effect (T={WT-cM, T-cM}) and S is the sample effect (S={sample1, sample2, sample3}). All raw p-values were corrected for multiple testing using the Benjamini-Hochberg (BH) method. Genes with an adjusted P <0.05 and a log2 fold-change >1 were called significant.

#### Pathway enrichment

We applied the pathway enrichment on the up-regulated genes (pvalue < 0.05 and logFC > 1) in T-cM and tumoroids versus to normal organoids by using KEGG, MSigDB curated gene sets and MSigDB regulatory target gene sets. The enrichment analysis was performed by R package clusterProfiler (v3.14.3) and msigdbr (v7.1.1).

Source code is available at this address: https://gist.github.com/wenjie1991/d79fe428ac80c8f2e5d781a966df3978

#### Gene set enrichment analysis

GSEA v4.0.3 was used to generate and calculate the enrichment score. Transcriptional signatures used for the analysis were extracted from the literature (Gregorieff *et al*, 2015; Yui *et al*, 2018; Merlos-Suárez *et al*, 2011; Mourao *et al*, 2019) and Nusse Lab (https://web.stanford.edu/group/nusselab/cgi-bin/wnt/target_genes). GSEA were calculated by gene set and 1000 permutations in our RNASeq normalised reads count matrix.

### Human colon cancer gene expression analysis

The gene expression data and clinical variables from the TCGA Colon Adenoma (COAD) cohort were downloaded from TSVdb (Sun *et al*, 2018) on 20th Jun 2020. Analyses were performed on Primary Solid Tumour gene expression data subset. Pairwise correlation analysis assayed THBS1, CTGF, CYR61 and WWC2 on 285 COAD tumour samples. The expression data were transformed by log2. Then, the Spearman correlation was calculated and visualized by the PerformanceAnalytics (v2.0.4) R package. In the survival analysis, 268 COAD tumour samples with overall survival follow-up information were used. The original data time scale in days was converted to years (365 days per year) and analysed up to 5 years. We defined the samples with upper one-third THBS1 expression as “THBS1 high group” and rests as “THBS1 low group”. Log-rank test and Kaplan Meier graph were obtained using the R package survminer (v0.4.7).

Source code is available at this address: https://gist.github.com/wenjie1991/6ff60b3edd5f61d0bd2ebe4f9404e46e

## Data and materials availability

The mass spectrometry proteomics data have been deposited to the ProteomeXchange Consortium via the PRIDE partner repository with the dataset identifier PXD020002(Perez-Riverol *et al*, 2019). The RNA sequencing data have been deposited in the Gene Expression Omnibus (GEO) repository under accession code GSE153160: Whole-genome transcriptomic analysis of intestinal organoids and tumoroids. All other data supporting the conclusions of this study are provided in the main text or the supplementary materials.

## Acknowledgments

We are very grateful to Prof. Shahragim Tajbakhsh for the mTmG reporter line, and to Prof. Hans Clevers for generously sharing the Lgr5-GFP (Barker *et al*, 2007) mice. We are also thankful to Dr. Jean-Leon Maitre, Dr. Raphael Margueron, and their team members (especially Dr. Daniel Holoch and Dr. Michel Wassef) for technical advice and constructive discussions. We would like to acknowledge the PICT-IBiSA imaging platform and the Flow Cytometry and Cell Sorting Platform at Institute Curie for their expertise; the In Vivo Experimental Facility, mainly Sonia Jannet, for help in the maintenance and care of our mouse colony as well as the NGS platform of Institut Curie for RNA sequencing and Dr. Nicolas Servant for bioinformatics support.

## Author contributions

G.J. and S.F. conceptualised and designed the project and wrote the manuscript S.F. also acquired funding and contributed to the data analysis. G.J. performed experiments, curated and interpreted the data, performed data visualisation, image analysis and quantification and prepared the figures. M.H. performed experiments and image analysis, analysed and interpreted data. W.S. curated the data and conducted formal data analysis. A.W., F.Q., M.P., Z.H., C.M., S.R. performed experiments. F.D. and G.A. performed and analysed, respectively, the mass Spectrometry data. D.L. supervised the Mass Spectrometry experiments. Z.H. and J.P. provided and induced the villinCre^ERT2^/Apc^flox/flox^ mouse model. D.V. provided the lifeact-GFP mouse model. S.F. provided funding, project administration and supervision.

## Materials & Correspondence

further information and requests for resources and reagents should be directed to and will be fulfilled by the corresponding author Silvia Fre.

---

**Table S1.**
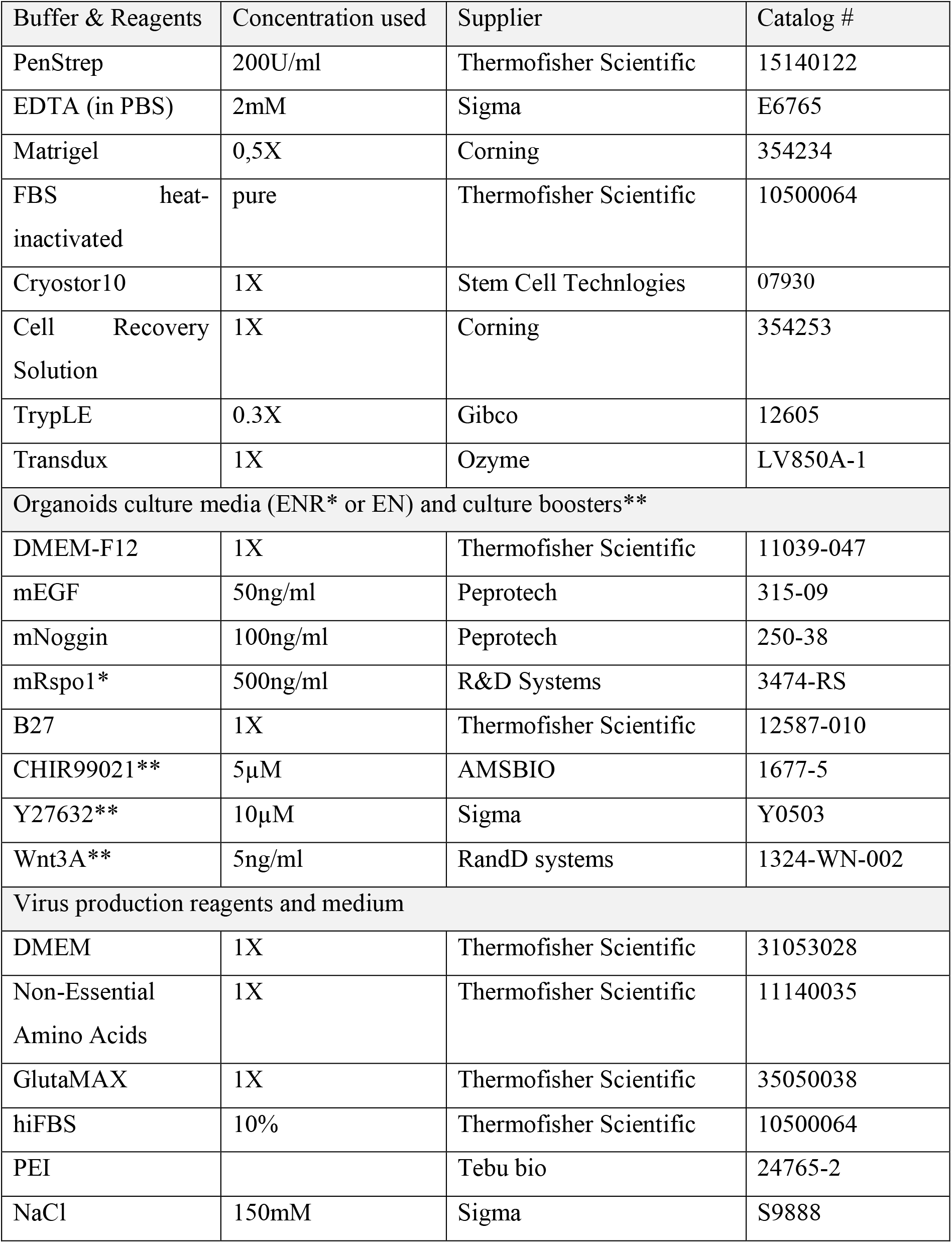
Culture reagents and buffer.

**Table S2.** Synthetic factors, inhibitors and neutralising antibodies.

| Components | Concentration used | Brand | Reference |
| --- | --- | --- | --- |
| Mouse anti-ANG | 2.5-25µg/ml | Abcam | Ab10600 |
| Rabbit anti-CP | 2.5-25µg/ml | Abcam | Ab48614 |
| Rabbit anti-CTGF | 1.25-12.5µg/ml | BioTechne | MAB91901-100 |
| Rabbit anti-HDGF | 0.5-5µg/ml | BioTechne | NBP1-71926 |
| Mouse anti-LGALS3 | 2.5-25µg/ml | Abcam | Ab2785 |
| Rabbit anti-LGALS3BP | 2.5-25µg/ml | Abcam | Ab217760 |
| Sheep anti-TTR | 63-120µg/ml | Abcam | Ab9015 |
| Mouse anti-THBS1 A6.1 | 5-20µg/ml | BioTechne | NB100-2059 |
| Mouse anti-THBS1 A4.1 | 5-20µg/ml | Thermofisher<br>scientific | MA5-13377 |
| Mouse anti-THBS1 C6.7 | 5-20µg/ml | Thermofisher<br>scientific | MA5-13390 |
| Mouse IgG1 Isotype<br>Control (MG1K) | 5-20µg/ml | BioTechne | NBP1-96983 |
| Mouse anti-FLAG | 1µg/ml | Sigma | F1804 |
| rmTHBS1 | 1-5µg/ml | BioTechne | 7859-TH-050 |
| Verteporfin | 5-10µM | Sigma | SML0534 |

**Table S3.**
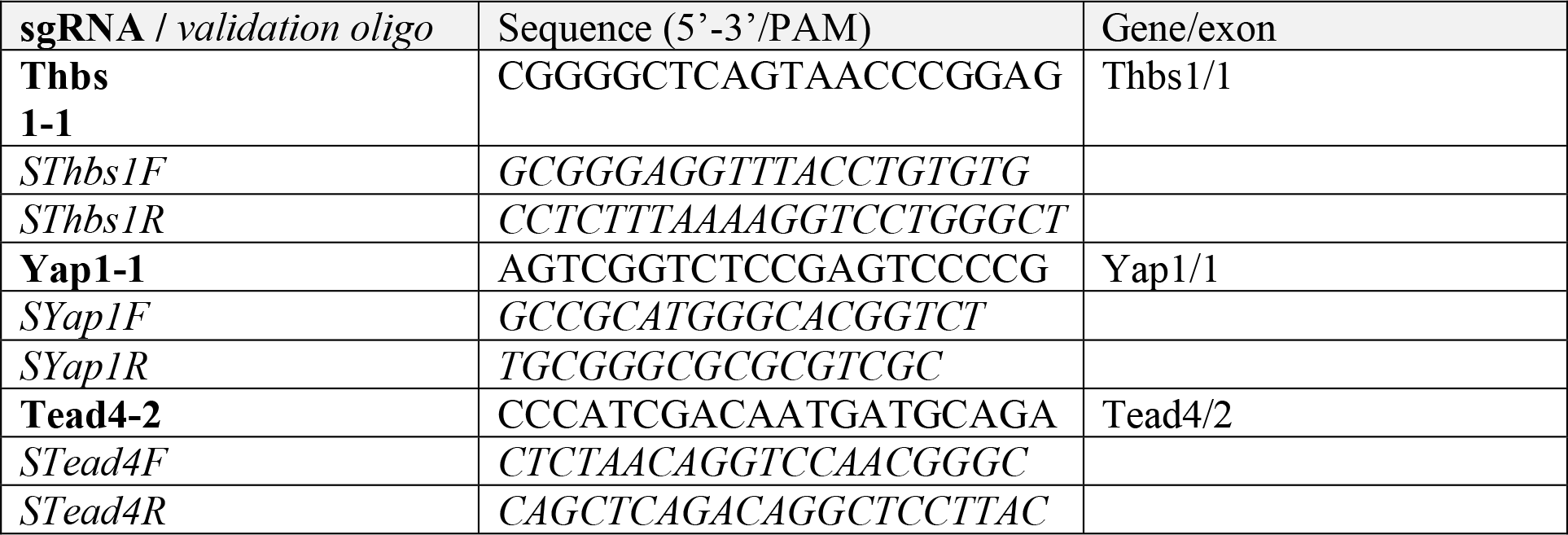
CRISPR sgRNA & validation oligos.

**Table S4.**
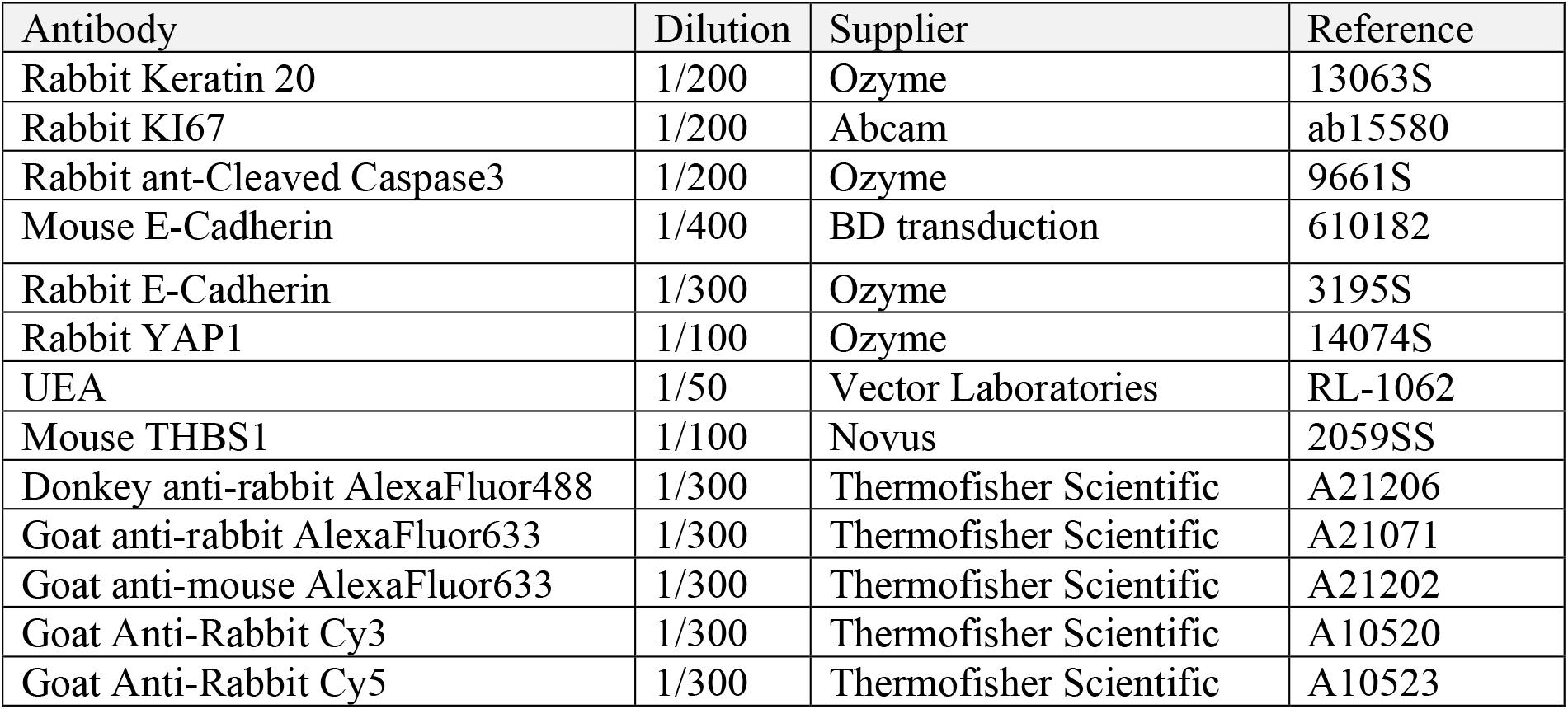
Antibodies.

| Antibody | Dilution | Supplier | Reference |
| --- | --- | --- | --- |
| Rabbit Keratin 20 | 1/200 | Ozyme | 13063S |
| Rabbit KI67 | 1/200 | Abcam | ab15580 |
| Rabbit ant-Cleaved Caspase3 | 1/200 | Ozyme | 9661S |
| Mouse E-Cadherin | 1/400 | BD transduction | 610182 |
| Rabbit E-Cadherin | 1/300 | Ozyme | 3195S |
| Rabbit YAP1 | 1/100 | Ozyme | 14074S |
| UEA | 1/50 | Vector Laboratories | RL-1062 |
| Mouse THBS1 | 1/100 | Novus | 2059SS |
| Donkey anti-rabbit AlexaFluor488 | 1/300 | Thermofisher Scientific | A21206 |
| Goat anti-rabbit AlexaFluor633 | 1/300 | Thermofisher Scientific | A21071 |
| Goat anti-mouse AlexaFluor633 | 1/300 | Thermofisher Scientific | A21202 |
| Goat Anti-Rabbit Cy3 | 1/300 | Thermofisher Scientific | A10520 |
| Goat Anti-Rabbit Cy5 | 1/300 | Thermofisher Scientific | A10523 |

**Fig. EV1.**
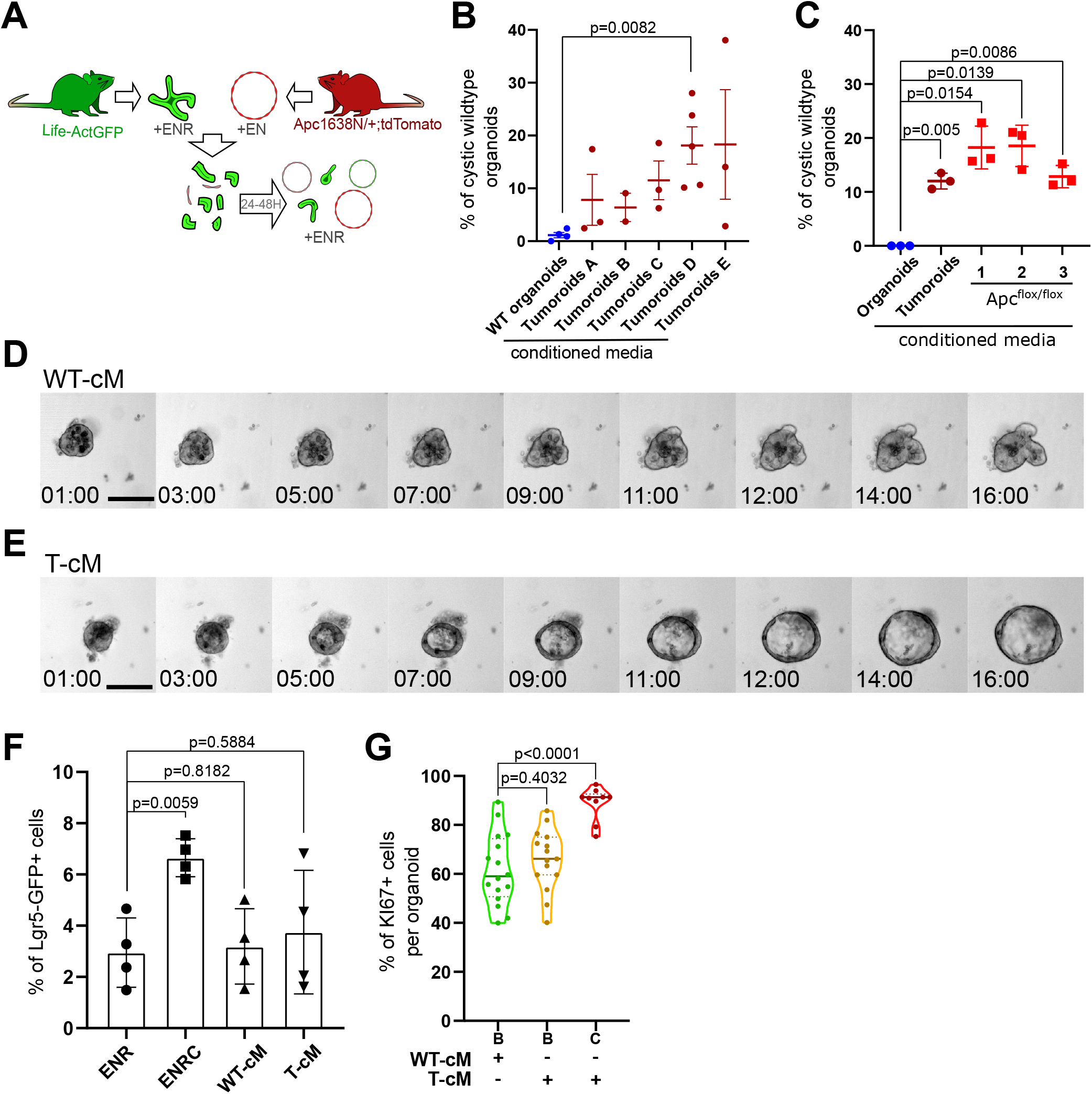
**Related to Figure 1.** (A) Experimental design of co-culture experiments with green organoids and red tumoroids at 3:1 ratio (WT organoids:tumoroids). ENR: medium containing EGF, Noggin and R-spondin1. (B) Quantification of the percentage of WT cystic organoids in the presence of cM derived from the indicated cultures. (C) Quantification of the percentage of WT cystic organoids in the presence of cM derived from WT organoids, tumoroids or 3 different Apc^-/-^ organoids generated from hyperplastic small intestine of VillinCre^ERT2^;Apc^flox/flox^ mice. (D-E) Time-lapse analysis of the growth of WT organoids in the presence of WT-cM (D) or T-cM (E) (in hours:minutes). (F) Quantification of the percentage of Lgr5-GFP+ cells in WT organoids treated with ENR, ENR + CHIR99021 (ENRC), WT-cM or T-cM (n=4 for each condition). (G) Quantification of the percentage of Ki67+ proliferative cells per organoid for budding organoids grown in WT-cM (B – WT-cM, n=16), budding organoids grown in T-cM (B – TcM, n=15) or cystic organoids grown in T-cM (C – T-cM, n=9). Scale bar = 100µm. Graphs indicate average values ± SD. Statistical analysis was performed with two-tailed unpaired Welch’s t-tests.

**Fig. EV2.**
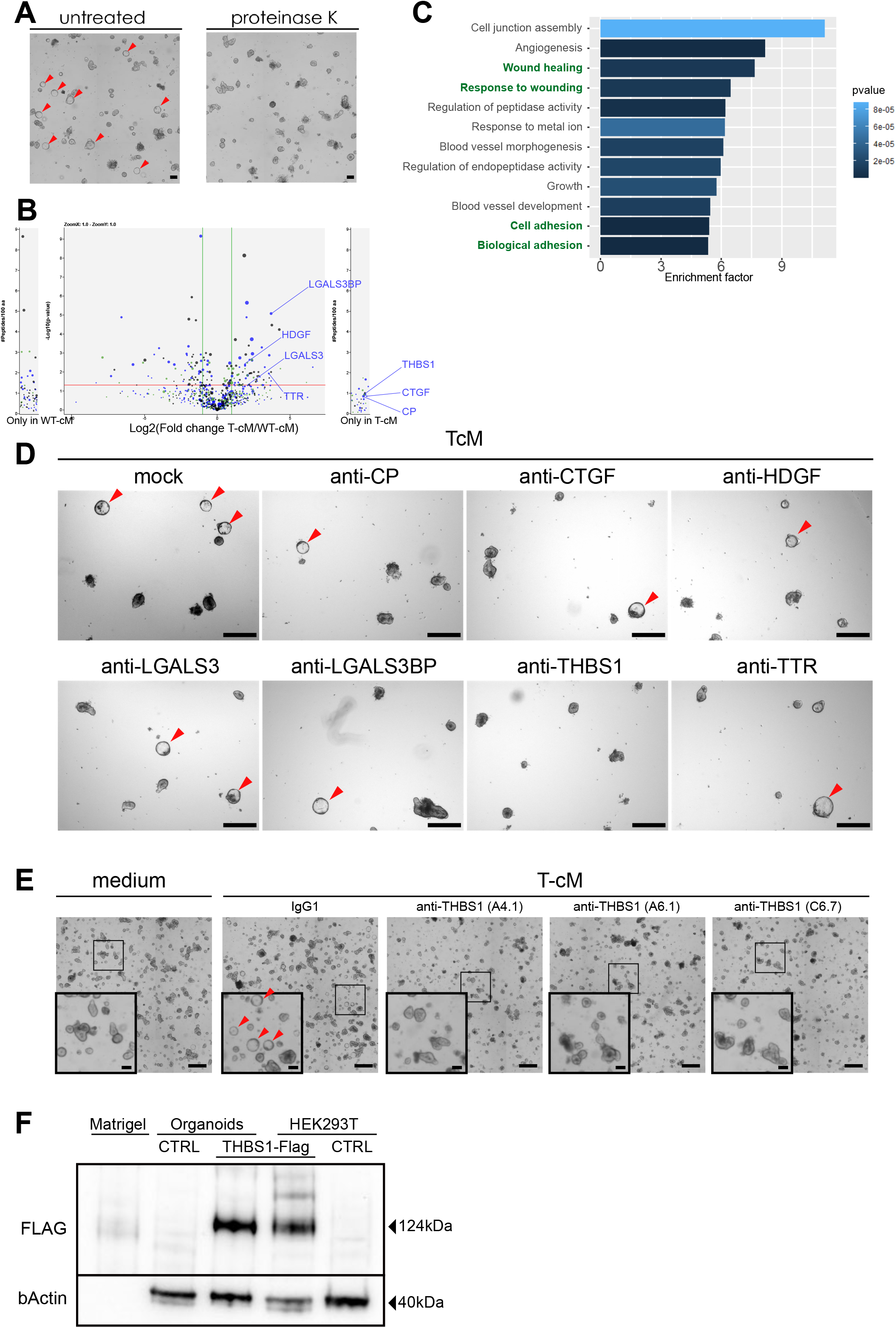
**Related to Figure 2.** (A) Proteinase K treatment of T-cM abolished the cystic transformation. (B) Volcano plot showing differential enrichment of proteins in T-cM compared to WT-cM by SILAC-based mass spectrometry. The selected candidates are indicated for one of the replicates in blue. The three different colours (black, blue and green dots) correspond to three replicates (n=3). (C) GO analysis of significantly enriched or unique proteins in T-cM compared to WT-cM. (D) Representative bright field images of WT organoids grown in T-cM upon neutralisation with blocking antibodies against the indicated proteins. (E) Representative bright field images of WT organoids grown in normal medium (left panel) or T-cM upon neutralisation with control IgG1 or three different blocking antibodies against THBS1 (clones A4.1, A6.1 and C6.7). (F) Western-blot anti-FLAG (targeting overexpressed THBS1-FLAG) and beta-actin (loading control) in: pure Matrigel, WT organoids infected with a control lentivirus (CTRL) or with lenti-Thbs1 (THBS1-Flag), HEK293T cells infected with lenti-Thbs1 (THBS1-Flag) or control lentivirus (CTRL). Scale bar = 100µm in A, 500µm in D-E (100µm in insets in E). Cystic organoids are indicated by red arrowheads in A, D and E.

**Fig. EV3.**
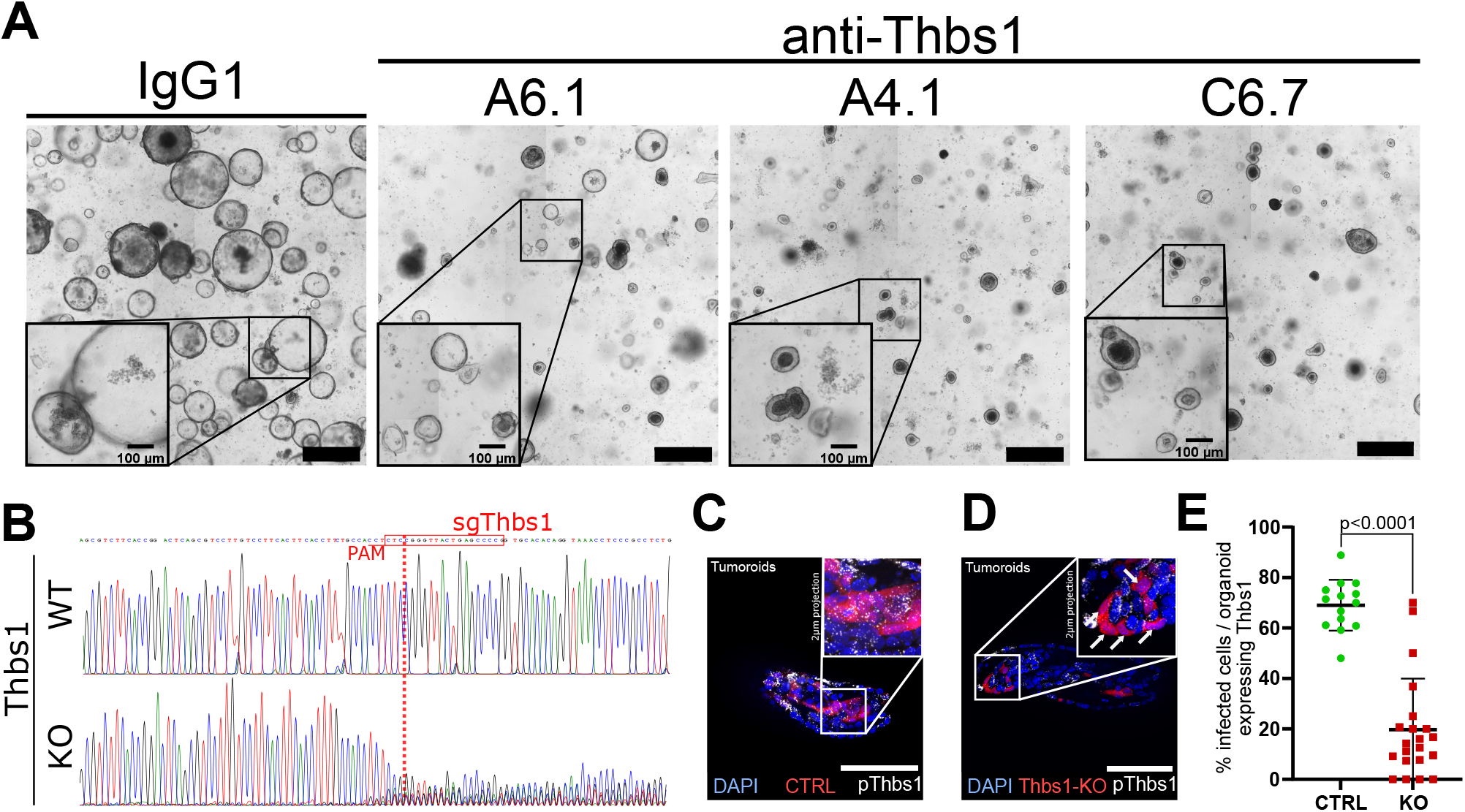
**Related to Figure 3.** (A) Representative images of tumoroids cultures grown for 48h in EN medium in the presence of control IgG1 or anti-THBS1 neutralising antibodies A6.1, A4.1, C6.7, as indicated. (B) Analysis of CRISPR edits from Sanger sequencing of DNA at the expected cut site for Cas9 in WT or THBS1 KO Mouse Embryonic Fibroblasts infected with no sgRNA or sgRNA targeting Thbs1, respectively. Red boxes indicate the sgRNA sequences with the PAM sequence underlined in red; red vertical dashed lines indicate the expected cut site. (C-D) Representative images of tumoroids infected with lenti-CRISPR expressing no sgRNA (CTRL in C) or sgThbs1 (KO in D) probed for Thbs1 expression using smRNA FISH with a probe recognising Thbs1 (pThbs1 in white). Red fluorescence indicates infected cells and DAPI stains the nuclei. (E) Quantification of the percentage of infected cells (red) expressing Thbs1 per organoid (n=3). Scale bar = 500µm in (A) and 100µm in C, D and insets in A. Statistical analysis was performed with two-tailed unpaired Welch’s t-tests.

**Fig. EV4.**
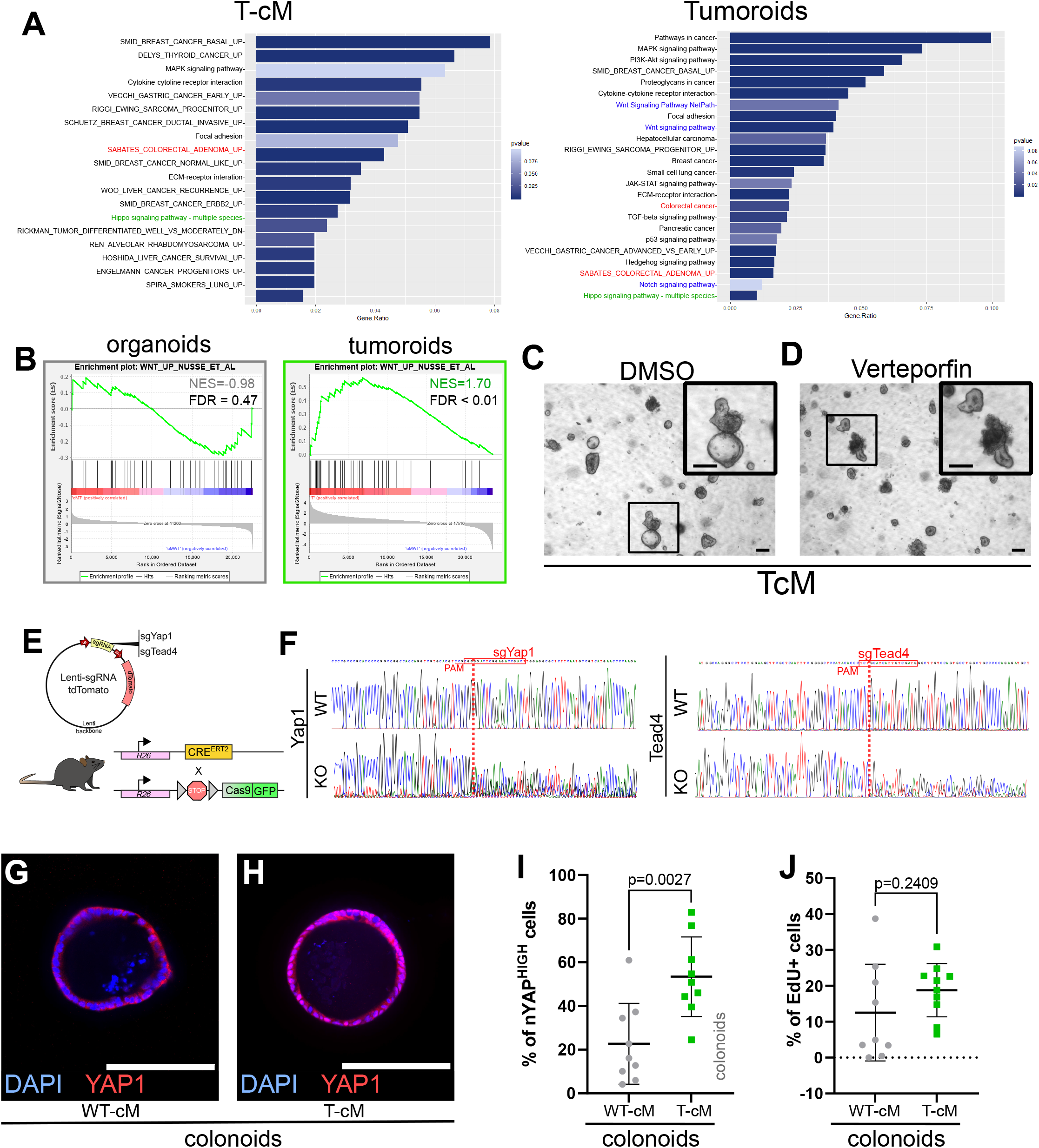
**Related to Figure 4.** (A) Pathway analysis of selected KEGG and GSEA-MSigDB terms enriched in the signatures of organoids grown in T-cM (left panel) or tumoroids (right panel). (B) Gene Set Enrichment Analysis (GSEA) comparing differentially expressed genes in WT organoids exposed to T-cM or in tumoroids with the WNT signature (Nusse Lab). Green NES = positive correlation; grey NES = non-significant correlation. (C-D) Representative bright field images of WT organoids grown in T-cM in the presence of DMSO (C) or the YAP inhibitor Verteporfin (D). (E) Map of the lenti- sgRNA-tdTomato lentiviral vector used for knock-out experiments and schematic diagram of the RosaCRE^ERT2^;Cas9-GFP mouse line used to derive WT organoids. (F) Analysis of CRISPR edits from Sanger sequencing of DNA at the expected cut site for Cas9 in Mouse Embryonic Fibroblasts infected with no sgRNA (WT) or sgRNAs targeting Yap1 (KO in F left) and Tead4 (KO in F right). Red boxes indicate the sgRNA sequences with the PAM sequence underlined in red; red vertical dashed lines indicate the expected cut site. (G-H) Representative immunofluorescence against YAP1 (in red) of colonoids exposed to WT-cM (G) or T-cM (H) for 24h. DAPI stains DNA in blue. (I-J) Quantification of the percentage of YAP^HIGH^ cells/organoid based on the ratio of nuclear vs. cytoplasmic YAP1 (I) or of EdU+ cells/organoid (J) in WT colonoids grown in WT- cM or T-cM. Scale bar = 100µm. Statistical analysis was performed with two-tailed unpaired Welch’s t-tests.

**Fig. EV5.**
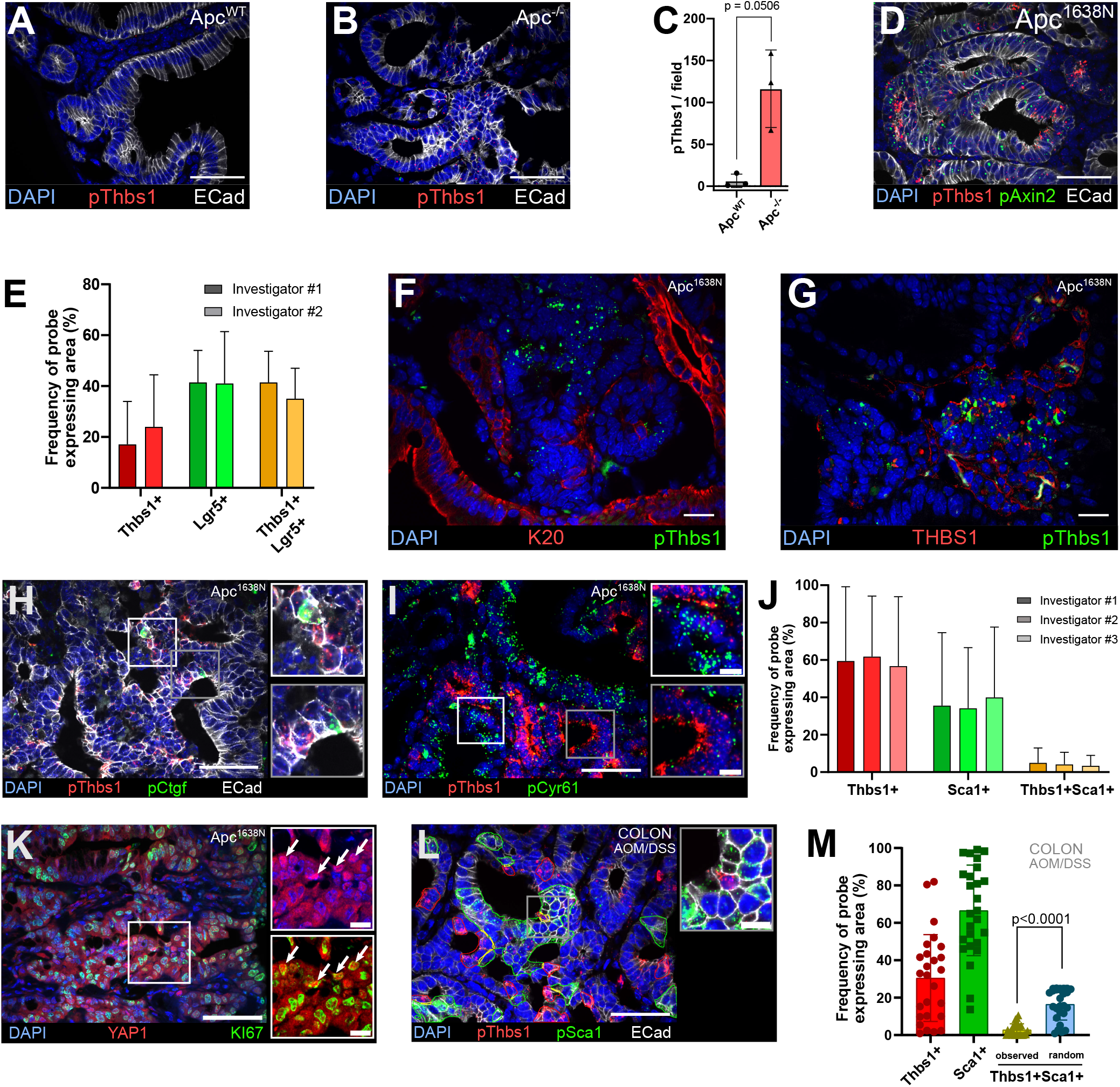
**Related to Figure 5.** (A-B) Representative sections of smRNA FISH targeting Thbs1 (pThbs1, red dots) in the small intestine of WT mice (Apc^WT^ in A) or villinCre^ERT2^;Apc^flox/flox^ mice (Apc^-/-^ in B). (C) Quantification of the number of Thbs1 dots per field in Apc^WT^ or Apc^-/-^ small intestine. (D) Representative sections of smRNA FISH targeting Thbs1 (pThbs1, red dots) and the Wnt target Axin2 (pAxin2, green dots) in mouse intestinal tumours (Apc^1638N^). (E) Quantification of the percentage of probe-expressing area reflecting cells expressing Thbs1 only (red), Lgr5 only (green) or co-expressing both (yellow) annotated by three independent and double-blinded investigators, corresponding to Fig. 5A-B. (F-G) Representative sections of mouse intestinal tumours analysed by smRNA FISH for Thbs1 (pThbs1, green dots) and immuno-stained with an antibody against the epithelial marker Keratin 20 (K20 in red in F) or with anti-THBS1 antibody (in red in G). (H-I) Representative sections of Apc mutant intestinal tumours analysed by smRNA FISH for the YAP target genes Ctgf (pCtgf, green dots in H) or Cyr61 (pCyr61, green dots in I) and Thbs1 (pThbs1, red dots). (J) Quantification of the percentage of probe-expressing area reflecting cells expressing Thbs1 only (red), Sca1 only (green) or co-expressing both (yellow) annotated by three independent and double-blinded investigators, corresponding to Fig. 5C-D. (K) Representative sections of Apc mutant intestinal tumours immuno-stained for YAP1 (in red) and KI67 (in green). White arrows in insets show nuclear YAP1^HIGH^ / Ki67+ cells (n=6 sections from 3 tumours). (L) Representative sections of chemically-induced colon tumours analysed by smRNA FISH for expression of Thbs1 (pThbs1, red dots) and the YAP target gene Sca1 (pSca1, green dots); regions presenting co-localisation of Thbs1 and Sca1 are outlined in yellow. (M) Quantification of the percentage of probe-expressing area reflecting cells expressing Thbs1 only (red), Sca1 only (green) or both (yellow). The observed mutual exclusion of red and green regions is statistically significant compared to the calculated probability of random co-expression (blue column) (n=30 sections from 5 tumours). E-cadherin antibody in white demarcates epithelial cells in A, B, D, H and L. DAPI labels nuclei in blue. Scale bar = 50µm (10µm in the insets). Statistical analysis was performed with two-tailed unpaired Welch’s t-tests in C (P=0.0506) and Wilcoxon test in M.

## References

Azzolin L, Panciera T, Soligo S, Enzo E, Bicciato S, Dupont S, Bresolin S, Frasson C, Basso G, Guzzardo V, et al (2014) YAP/TAZ incorporation in the β-catenin destruction complex orchestrates the Wnt response. Cell 158: 157–170

Balkwill FR, Capasso M & Hagemann T (2012) The tumor microenvironment at a glance. J Cell Sci 125: 5591–5596

Barker N, van Es JH, Kuipers J, Kujala P, van den Born M, Cozijnsen M, Haegebarth A, Korving J, Begthel H, Peters PJ, et al (2007) Identification of stem cells in small intestine and colon by marker gene Lgr5. Nature 449: 1003–1007

Barker N, Ridgway RA, van Es JH, van de Wetering M, Begthel H, van den Born M, Danenberg E, Clarke AR, Sansom OJ & Clevers H (2009) Crypt stem cells as the cells-of-origin of intestinal cancer. Nature 457: 608–611

Barry ER, Morikawa T, Butler BL, Shrestha K, de la Rosa R, Yan KS, Fuchs CS, Magness ST, Smits R, Ogino S, et al (2013) Restriction of intestinal stem cell expansion and the regenerative response by YAP. Nature 493: 106–110

Benjamini Y & Hochberg Y (1995) Controlling the False Discovery Rate: A Practical and Powerful Approach to Multiple Testing. Journal of the Royal Statistical Society Series B (Methodological) 57: 289–300

Brugmann SA, Goodnough LH, Gregorieff A, Leucht P, ten Berge D, Fuerer C, Clevers H, Nusse R & Helms JA (2007) Wnt signaling mediates regional specification in the vertebrate face. Development 134: 3283–3295

Cai J, Maitra A, Anders RA, Taketo MM & Pan D (2015) β-Catenin destruction complex-independent regulation of Hippo–YAP signaling by APC in intestinal tumorigenesis. Genes & Development 29: 1493–1506

Cheung P, Xiol J, Dill MT, Yuan W-C, Panero R, Roper J, Osorio FG, Maglic D, Li Q, Gurung B, et al (2020) Regenerative Reprogramming of the Intestinal Stem Cell State via Hippo Signaling Suppresses Metastatic Colorectal Cancer. Cell Stem Cell 27: 590–604.e9

Cleary AS, Leonard TL, Gestl SA & Gunther EJ (2014) Tumour cell heterogeneity maintained by cooperating subclones in Wnt-driven mammary cancers. Nature 508: 113–117

Colnot S, Decaens T, Niwa-Kawakita M, Godard C, Hamard G, Kahn A, Giovannini M & Perret C (2004) Liver-targeted disruption of Apc in mice activates -catenin signaling and leads to hepatocellular carcinomas. Proceedings of the National Academy of Sciences 101: 17216–17221

Dobin A, Davis CA, Schlesinger F, Drenkow J, Zaleski C, Jha S, Batut P, Chaisson M & Gingeras TR (2013) STAR: ultrafast universal RNA-seq aligner. Bioinformatics 29: 15–21

Drost J, van Jaarsveld RH, Ponsioen B, Zimberlin C, van Boxtel R, Buijs A, Sachs N, Overmeer RM, Offerhaus GJ, Begthel H, et al (2015) Sequential cancer mutations in cultured human intestinal stem cells. Nature 521: 43–47

Flanagan DJ, Pentinmikko N, Luopajärvi K, Willis NJ, Gilroy K, Raven AP, Mcgarry L, Englund JI, Webb AT, Scharaw S, et al (2021) NOTUM from Apc-mutant cells biases clonal competition to initiate cancer. Nature 594: 430–435

Fodde R, Edelmann W, Yang K, van Leeuwen C, Carlson C, Renault B, Breukel C, Alt E, Lipkin M & Khan PM (1994) A targeted chain-termination mutation in the mouse Apc gene results in multiple intestinal tumors. Proc Natl Acad Sci USA 91: 8969–8973

Germann M, Xu H, Malaterre J, Sampurno S, Huyghe M, Cheasley D, Fre S & Ramsay RG (2014) Tripartite interactions between Wnt signaling, Notch and Myb for stem/progenitor cell functions during intestinal tumorigenesis. Stem Cell Research 13: 355–366

Gregorieff A, Liu Y, Inanlou MR, Khomchuk Y & Wrana JL (2015) Yap-dependent reprogramming of Lgr5(+) stem cells drives intestinal regeneration and cancer. Nature 526: 715–718

Guillermin O, Angelis N, Sidor CM, Ridgway R, Baulies A, Kucharska A, Antas P, Rose MR, Cordero J, Sansom O, et al (2021) Wnt and Src signals converge on YAP-TEAD to drive intestinal regeneration. EMBO J

Gutierrez LS, Suckow M, Lawler J, Ploplis VA & Castellino FJ (2003) Thrombospondin 1--a regulator of adenoma growth and carcinoma progression in the APC(Min/+) mouse model. Carcinogenesis 24: 199–207

Hanahan D & Coussens LM (2012) Accessories to the Crime: Functions of Cells Recruited to the Tumor Microenvironment. Cancer Cell 21: 309–322

Jardé T, Evans RJ, McQuillan KL, Parry L, Feng GJ, Alvares B, Clarke AR & Dale TC (2013) In vivo and in vitro models for the therapeutic targeting of Wnt signaling using a Tet-OΔN89β- catenin system. Oncogene 32: 883–893

Krotenberg Garcia A, Fumagalli A, Le HQ, Jackstadt R, Lannagan TRM, Sansom OJ, van Rheenen J & Suijkerbuijk SJE (2021) Active elimination of intestinal cells drives oncogenic growth in organoids. Cell Reports 36: 109307

Liu-Chittenden Y, Huang B, Shim JS, Chen Q, Lee S-J, Anders RA, Liu JO & Pan D (2012) Genetic and pharmacological disruption of the TEAD-YAP complex suppresses the oncogenic activity of YAP. Genes & Development 26: 1300–1305

Lopez-Dee ZP, Chittur SV, Patel H, Chinikaylo A, Lippert B, Patel B, Lawler J & Gutierrez LS (2015) Thrombospondin-1 in a Murine Model of Colorectal Carcinogenesis. PLoS ONE 10: e0139918

el Marjou F, Janssen K-P, Chang BH-J, Li M, Hindie V, Chan L, Louvard D, Chambon P, Metzger D & Robine S (2004) Tissue-specific and inducible Cre-mediated recombination in the gut epithelium. Genesis 39: 186–193

Marusyk A, Tabassum DP, Altrock PM, Almendro V, Michor F & Polyak K (2014) Non-cell- autonomous driving of tumour growth supports sub-clonal heterogeneity. Nature 514: 54– 58

Merlos-Suárez A, Barriga FM, Jung P, Iglesias M, Céspedes MV, Rossell D, Sevillano M, Hernando-Momblona X, da Silva-Diz V, Muñoz P, et al (2011) The Intestinal Stem Cell Signature Identifies Colorectal Cancer Stem Cells and Predicts Disease Relapse. Cell Stem Cell 8: 511–524

Mourao L, Jacquemin G, Huyghe M, Nawrocki WJ, Menssouri N, Servant N & Fre S (2019) Lineage tracing of Notch1-expressing cells in intestinal tumours reveals a distinct population of cancer stem cells. Sci Rep 9: 888

Muzumdar MD, Tasic B, Miyamichi K, Li L & Luo L (2007) A global double-fluorescent Cre reporter mouse. Genesis 45: 593–605

van Neerven SM, de Groot NE, Nijman LE, Scicluna BP, van Driel MS, Lecca MC, Warmerdam DO, Kakkar V, Moreno LF, Vieira Braga FA, et al (2021) Apc-mutant cells act as supercompetitors in intestinal tumour initiation. Nature 594: 436–441

Ombrato L, Nolan E, Kurelac I, Mavousian A, Bridgeman VL, Heinze I, Chakravarty P, Horswell S, Gonzalez-Gualda E, Matacchione G, et al (2019) Metastatic-niche labelling reveals parenchymal cells with stem features. Nature 572: 603–608

Ong S-E, Blagoev B, Kratchmarova I, Kristensen DB, Steen H, Pandey A & Mann M (2002) Stable Isotope Labeling by Amino Acids in Cell Culture, SILAC, as a Simple and Accurate Approach to Expression Proteomics. Mol Cell Proteomics 1: 376–386

Onuma K, Ochiai M, Orihashi K, Takahashi M, Imai T, Nakagama H & Hippo Y (2013) Genetic reconstitution of tumorigenesis in primary intestinal cells. Proceedings of the National Academy of Sciences 110: 11127–11132

Perez-Riverol Y, Csordas A, Bai J, Bernal-Llinares M, Hewapathirana S, Kundu DJ, Inuganti A, Griss J, Mayer G, Eisenacher M, et al (2019) The PRIDE database and related tools and resources in 2019: improving support for quantification data. Nucleic Acids Res 47: D442– D450

Platt RJ, Chen S, Zhou Y, Yim MJ, Swiech L, Kempton HR, Dahlman JE, Parnas O, Eisenhaure TM, Jovanovic M, et al (2014) CRISPR-Cas9 knockin mice for genome editing and cancer modeling. Cell 159: 440–455

Poullet P, Carpentier S & Barillot E (2007) myProMS, a web server for management and validation of mass spectrometry-based proteomic data. Proteomics 7: 2553–2556

Resovi A, Pinessi D, Chiorino G & Taraboletti G (2014) Current understanding of the thrombospondin-1 interactome. Matrix Biol 37: 83–91

Riedl J, Crevenna AH, Kessenbrock K, Yu JH, Neukirchen D, Bista M, Bradke F, Jenne D, Holak TA, Werb Z, et al (2008) Lifeact: a versatile marker to visualize F-actin. Nat Methods 5: 605–607

Ring DB, Johnson KW, Henriksen EJ, Nuss JM, Goff D, Kinnick TR, Ma ST, Reeder JW, Samuels I, Slabiak T, et al (2003) Selective glycogen synthase kinase 3 inhibitors potentiate insulin activation of glucose transport and utilization in vitro and in vivo. Diabetes 52: 588–595

Ritchie ME, Phipson B, Wu D, Hu Y, Law CW, Shi W & Smyth GK (2015) limma powers differential expression analyses for RNA-sequencing and microarray studies. Nucleic Acids Res 43: e47

Robinson MD, McCarthy DJ & Smyth GK (2010) edgeR: a Bioconductor package for differential expression analysis of digital gene expression data. Bioinformatics 26: 139–140

Sanjana NE, Shalem O & Zhang F (2014) Improved vectors and genome-wide libraries for CRISPR screening. Nat Methods 11: 783–784

Sato T & Clevers H (2013) Growing Self-Organizing Mini-Guts from a Single Intestinal Stem Cell: Mechanism and Applications. Science 340: 1190–1194

Sato T, Stange DE, Ferrante M, Vries RGJ, Van Es JH, Van den Brink S, Van Houdt WJ, Pronk A, Van Gorp J, Siersema PD, et al (2011) Long-term expansion of epithelial organoids from human colon, adenoma, adenocarcinoma, and Barrett’s epithelium. Gastroenterology 141: 1762–1772

Sato T, Vries RG, Snippert HJ, van de Wetering M, Barker N, Stange DE, van Es JH, Abo A, Kujala P, Peters PJ, et al (2009) Single Lgr5 stem cells build crypt-villus structures in vitro without a mesenchymal niche. Nature 459: 262–265

Schwank G, Koo B-K, Sasselli V, Dekkers JF, Heo I, Demircan T, Sasaki N, Boymans S, Cuppen E, van der Ent CK, et al (2013) Functional repair of CFTR by CRISPR/Cas9 in intestinal stem cell organoids of cystic fibrosis patients. Cell Stem Cell 13: 653–658

Shalem O, Sanjana NE, Hartenian E, Shi X, Scott DA, Mikkelsen TS, Heckl D, Ebert BL, Root DE, Doench JG, et al (2014) Genome-Scale CRISPR-Cas9 Knockout Screening in Human Cells. Science 343: 84–87

Sid B, Langlois B, Sartelet H, Bellon G, Dedieu S & Martiny L (2008) Thrombospondin-1 enhances human thyroid carcinoma cell invasion through urokinase activity. Int J Biochem Cell Biol 40: 1890–1900

Sun W, Duan T, Ye P, Chen K, Zhang G, Lai M & Zhang H (2018) TSVdb: a web-tool for TCGA splicing variants analysis. BMC Genomics 19: 405

Tanaka T, Kohno H, Suzuki R, Yamada Y, Sugie S & Mori H (2003) A novel inflammation-related mouse colon carcinogenesis model induced by azoxymethane and dextran sodium sulfate. Cancer Sci 94: 965–973

Taniguchi K, Moroishi T, de Jong PR, Krawczyk M, Grebbin BM, Luo H, Xu R, Golob-Schwarzl N, Schweiger C, Wang K, et al (2017) YAP–IL-6ST autoregulatory loop activated on APC loss controls colonic tumorigenesis. Proc Natl Acad Sci USA 114: 1643–1648

Taniguchi K, Wu L-W, Grivennikov SI, de Jong PR, Lian I, Yu F-X, Wang K, Ho SB, Boland BS, Chang JT, et al (2015) A gp130-Src-YAP module links inflammation to epithelial regeneration. Nature 519: 57–62

Tuszynski GP, Gasic TB, Rothman VL, Knudsen KA & Gasic GJ (1987) Thrombospondin, a potentiator of tumor cell metastasis. Cancer Res 47: 4130–4133

Ventura A, Kirsch DG, McLaughlin ME, Tuveson DA, Grimm J, Lintault L, Newman J, Reczek EE, Weissleder R & Jacks T (2007) Restoration of p53 function leads to tumour regression in vivo. Nature 445: 661–665

Yamashiro Y, Thang BQ, Ramirez K, Shin SJ, Kohata T, Ohata S, Nguyen TAV, Ohtsuki S, Nagayama K & Yanagisawa H (2019) Matrix mechanotransduction mediated by thrombospondin-1/integrin/YAP signaling pathway in the remodeling of blood vessels Molecular Biology

Yui S, Azzolin L, Maimets M, Pedersen MT, Fordham RP, Hansen SL, Larsen HL, Guiu J, Alves MRP, Rundsten CF, et al (2018) YAP/TAZ-Dependent Reprogramming of Colonic Epithelium Links ECM Remodeling to Tissue Regeneration. Cell Stem Cell 22: 35–49.e7

Yum MK, Han S, Fink J, Wu S-HS, Dabrowska C, Trendafilova T, Mustata R, Chatzeli L, Azzarelli R, Pshenichnaya I, et al (2021) Tracing oncogene-driven remodelling of the intestinal stem cell niche. Nature 594: 442–447

Zanconato F, Cordenonsi M & Piccolo S (2016) YAP/TAZ at the Roots of Cancer. Cancer Cell 29: 783–803

